# Machine learning analysis reveals the dynamics of mode transition in dendritic cell migration

**DOI:** 10.1101/2022.07.07.499070

**Authors:** Taegeun Song, Youngjun Choi, Jae-Hyung Jeon, Yoon-Kyoung Cho

**Author notes:** Correspondence and requests for materials: Addressed to J.H. J. or Y-K. C. These authors contributed equally to this work (Taegeun Song, Youngjun Choi).

## Abstract

Dendritic cells (DCs) patrol the body as immunological sentinels and search for pathogens. Upon stimulation, immature DCs (imDCs) become mature DCs (mDCs), which migrate to the lymph nodes and present antigens to T cells. The migratory behavior is crucial for initiating and controlling immune responses; however, the properties of the highly heterogeneous and dynamic motility phenotype are not fully understood. Here, we established an unsupervised machine learning (ML) strategy to investigate spatiotemporal motility patterns in long-term, two-dimensional cell migration trajectories, and determined the number of motility patterns and how these are related to the maturation status. We identified three distinct migratory modes independent of the cell state: slow-diffusive (SD), slow-persistent (SP), and fast-persistent (FP). We found that maturation-dependent motility changes are emergent properties of the distribution and dynamic transitions of these three modes. Remarkably, imDCs changed their migration modes more frequently, and predominantly followed the SD→FP→SP→SD unicyclic transition, indicating that imDCs rapidly increase their speed during the shift from diffusive to persistent motility; however, persistence progressively declines when switching back to diffusive motility. In contrast, mDCs show no transition directionality. Our ML-promoted motility pattern analysis and history-dependent mode transition investigation may provide new insights into the complex process of biological motility.

---

Cell migration is essential for homeostasis in living systems^1^. Intriguingly, cell motility shows complex dynamics beyond the classical diffusion theory^2^. Therefore, various random-walk models have been employed to explain anomalous diffusion processes^3,4^. For example, bacterial micro swimmers and T cells use an intermittent search process as an effective search mechanism, which alternates between two types of slow/fast motion, such as the run-and-tumble motion and Lévy walk^5,6^.

Dendritic cells (DCs) exhibit adaptive motility patterns, reflecting their immunological function as major antigen-presenting cells^7,8^. Immature DCs (imDCs) typically navigate using an intermittent search strategy, which combines fast persistent motility during patrolling with slow diffusive motility for antigen collection^9–11^. In contrast, mature DCs (mDCs) mainly employ fast persistent motility, enabling them to reach lymph nodes and deliver antigens to T cells^12,13^. Recent single-cell studies employing microfabricated devices showed that deterministic actin waves contribute to the intermittent search mechanism^14^. Additionally, Lévy walk patterns were identified and showed directional persistence and zigzag motion as an *in vivo* search strategy^15^. Although extensive studies have been performed on the overall cellular migration characteristics and physiological framework of DC motility, the description of the average migration dynamics is mostly restricted to the dichotomous approach, such as slow or fast, diffusive or persistent, zigzag or non-zigzag.

We hypothesized that such simple interpretations overlook the complex and heterogeneous dynamics of single-cell motility. Intrigued by experimental observations of heterogeneous distribution of DC motility (**Extended data Fig. 1**), we designed an unsupervised machine learning (ML) method to uncover DC motility patterns at the single-cell level, and understand the distinct dynamic modes and their transition dynamics quantitatively. We used primary mouse bone-marrow-derived DCs (BMDCs) (**Supplementary Section A, Fig. S1**). The use of a gel confiner^13,16^ enabled long-term (24 h) monitoring of an unbiased population of freely migrating cells with precise control over the degree of confinement (**Fig. 1a, b, Supplementary Section B, Fig. S2**). Cellular movements were imaged every 1 min using brightfield microscopy with a millimeter-scale broad field of view (1.3 × 1.3 mm^2^) (**Supplementary Section C, Videos 1, 2**). Using live-cell tracking data (**Fig. 1c**), we found that mDCs showed faster and more persistent motility than imDCs, which is consistent with the results of previous studies (**Fig. 1d, e**)^12,13^. However, outliers were always present (**Fig. 1f**), and these atypical phenomena were often neglected in the pooled population analysis (**Extended data Fig. 1**).

**Fig. 1.**
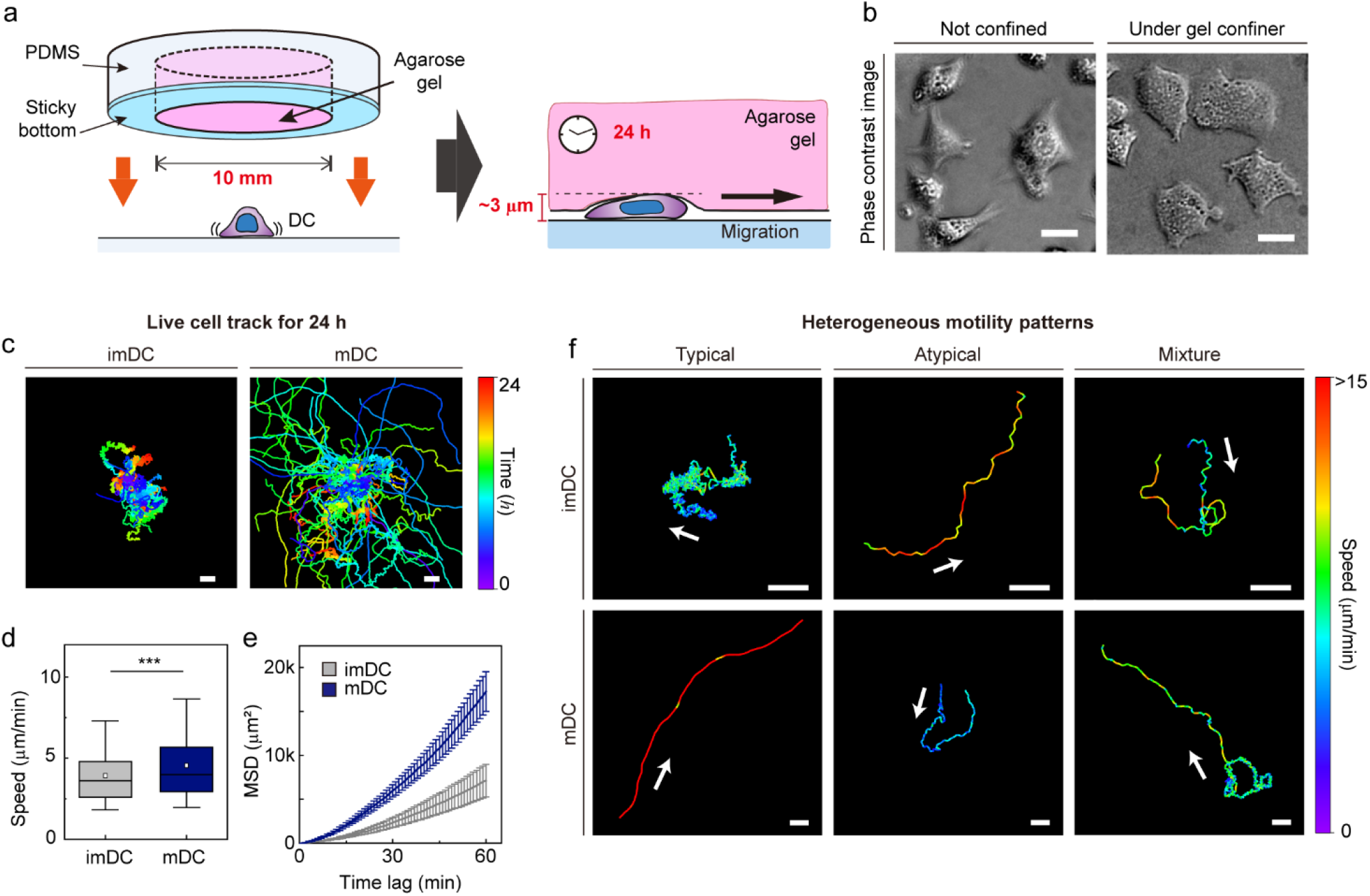
Long-term analysis of dendritic cell (DC) motility under confinement. **a**, Scheme of a gel confiner for an under-agarose assay with stable and reproducible confinement. The agarose gel (Young’s elastic modulus: 11.13 kPa) provided a tissue-like environment, and the sticky bottom beneath the PDMS structure enabled stable and reproducible confinement over a large area (78.5 mm^2^) **b**, Phase-contrast images of DCs with and without the gel confiner. Scale bar = 10 μm. **c**, 2D cellular migration trajectories of imDCs and mDCs for 24 h. The starting point of each trajectory was translated to the origin of the plot. Color codes indicate the track duration. Scale bar = 100 μm. One representative experiment out of three is shown. **d**, Mean track speed of imDCs and mDCs. In the box plots, the bars include 95% of the points, the center corresponds to the median, and the box contains 75% of the data. Data were pooled from three independent experiments (imDC n= 460; mDC n= 371). The Mann-Whitney test was used for comparing two groups. ***: *P* < 0.001. **e**, Mean square displacement (MSD) curves of imDCs and mDCs. Lines indicate averaged MSD from three independent experiments, and the error bar indicates the S.E. **f**, Examples of heterogeneous motility patterns of DCs. Motility directions are marked with arrows. The color codes indicate instantaneous speed. Scale bar = 100 μm.

ML analysis is a powerful tool for classifying complex natural phenomena. In the field of single-particle trajectory analysis^17^, ML has been exploited for reusing imperfect datasets^18^, retrieving information on location or polarization^19^, or inferring transport models^20,21^. In this study, we developed an ML method to quantitatively analyze complex and heterogeneous cellular motility processes. The cell migration trajectories were first segmented into 1 h long tracks. Subsequently, five features quantifying motility (radius of gyration *R*_*g*_, end-to-end distance *R*_*ete*_, and average kinetic energy *E*) and directionality (asphericity *A* and turning angle fluctuation Δ*θ*) were evaluated as the input bases for the machine kernel (**Fig. 2, Supplementary Section D**).

**Fig. 2.**
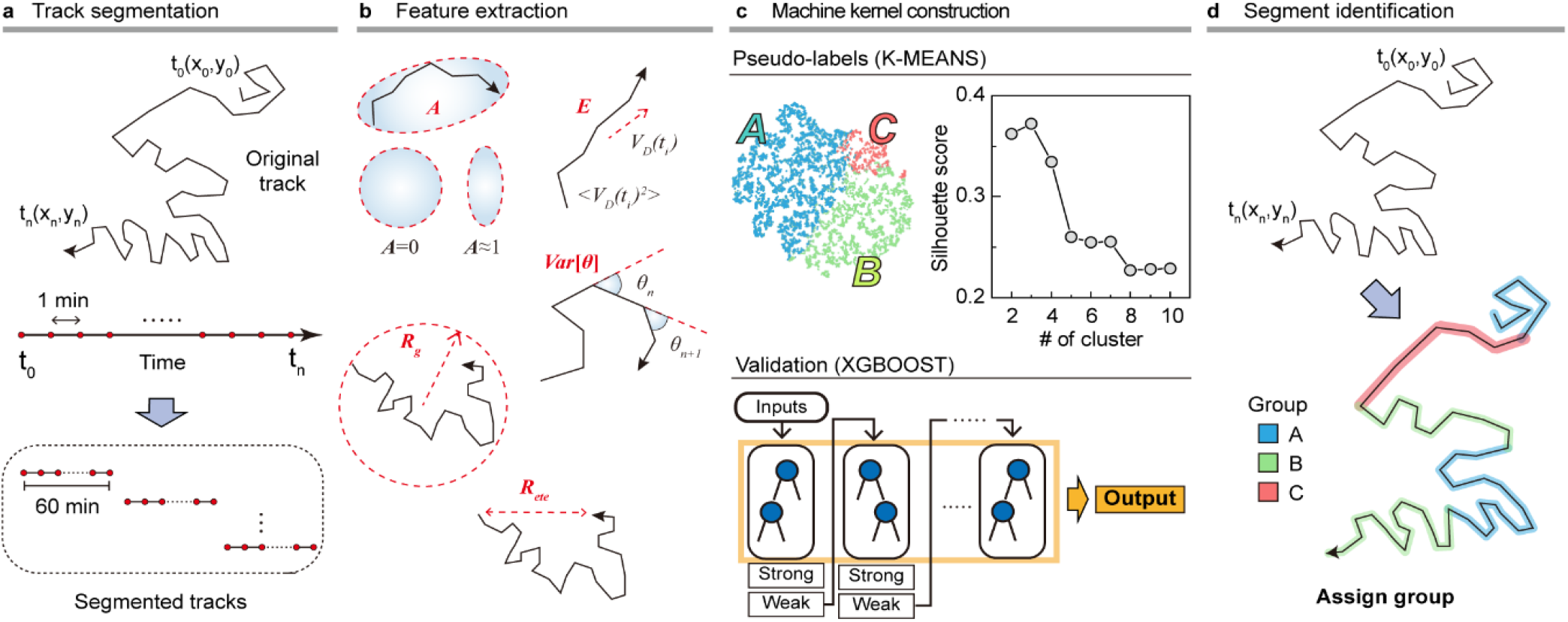
Overview of the machine learning method. **a**, Track segmentation. We obtained 649 raw trajectories from the training datasets. After performing the data pre-processing, we segmented the trajectory into 1 h long pieces indexed with their start and end times. Using 5741 segmented trajectories, we made a trajectory pool. **b**, Feature extraction. Using the segmented tracks, we calculated five features: radius of gyration (*R*_*g*_), asphericity (*A*), energy consumption (*E*), end-to-end distance (*R*_*ete*_), and variance of turning angles (*Var*[*θ*]). **c**, The construction of the machine kernel. Using the input data, we performed K-means^37,38^ clustering to classify the trajectories using an unsupervised learning method and specify the label. Next, with the randomly chosen 2000 labeled trajectories, we trained XGBOOST^39,40^ using supervised learning. The trained XGBOOST showed 99.6% agreement with the results of K-means clustering for the label prediction. The hyperparameters were optimized using a cross-validated grid search in both methods. **d**, Segment identification. Using the trained XGBOOST, we analyzed the original trajectory with a 1 h time window to identify the dynamic state of each segment

Using K-means unsupervised clustering^22,23^, we found that the input data were classified into three groups (**Fig. 3a, b, Supplementary Section E, Extended data Fig. 2, 3**). Furthermore, the trained XGBOOST machine inferred that DC migration is classified into three distinct patterns: SD, FP, and SP modes (**Table 1, Supplementary videos 3–5**). DCs in the SD mode perform anti-persistent walks at a slow speed, whereas FP mode cells show directional random walks at a fast speed. The SP mode differed from the other two modes. It had a directional walk analogous to that of FP mode cells. However, the feature properties differed significantly from each other (**Extended data Fig. 3**). In terms of average speed, the SP mode was similar to the SD mode (**Extended data Fig. 3**).

**Table 1.** Summary of the characteristics of each group (dynamic mode).

|  | Group I | Group II | Group III |
| --- | --- | --- | --- |
| Average speed (µm/min) | 2.58 ± 1.90 | 2.21 ± 1.61 | 6.27 ± 2.66 |
| Anomalous exponent (*) | 0.59 | 1.47 | 1.67 |
| Directionality (*) | X | O | O |
| Dynamic mode | Slow-Diffusive (SD) | Slow-Persist (SP) | Fast-Persist (FP) |

(\*) is defined in **Supplementary section F**.

**Fig. 3.**
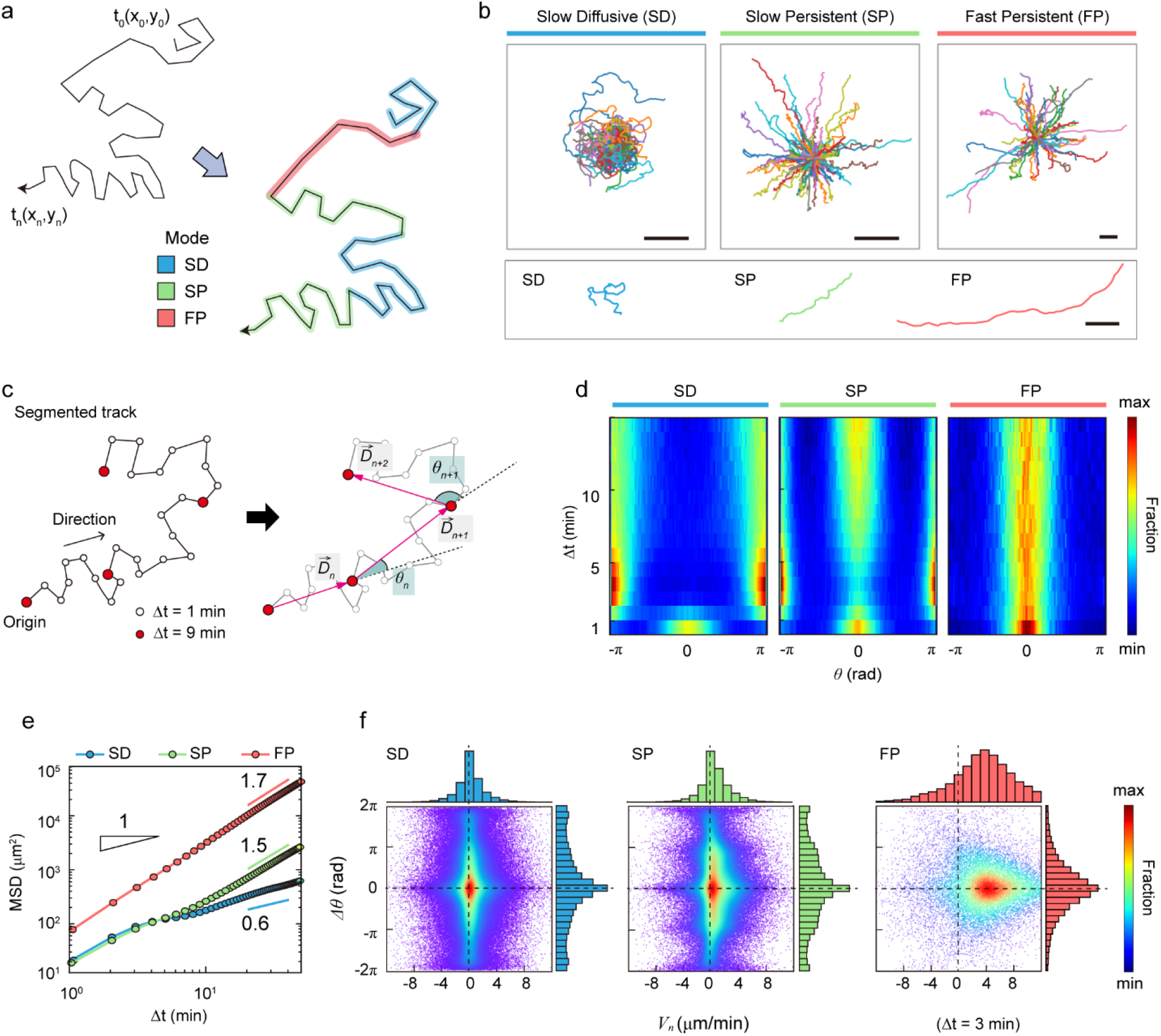
Statistical characteristics of the machine defined three distinct modes of DCs motility. **a**, Scheme of three distinct modes in a trajectory. **b**, Sample trajectories of slow diffusive (SD), slow persistent (SP), and fast persistent (FP) migration modes. Representative trajectories of each migration mode are shown together at the bottom panel (scale bar: 100 μm). **c**, Schematic trajectories with the definition of the displacement vector 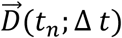 and turning angle *θ*_*n*_. The turning angle ranges from −*π* (clockwise turning) to *π* (counter-clock-wise turning). In the phase-space density map, we measured the magnitude difference for successive displacement vectors 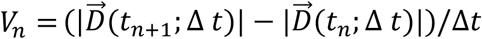 and their turning angle difference Δ*θ*_*n*_ = *θ*_*n*+1_ − *θ*_*n*_. **d**, The turning angle heat map (examples of individual trajectories are shown in **Extended data Fig. 6**). See its mathematical definition in **Supplementary Section F. e**, The MSD curves for SD (blue), SP (green), and FP (red) modes. The guidelines show the anomalous exponent at long times. **f**, The density map of phase space (*V*_*n*_, Δ*θ*_*n*_) at Δ*t* = 3 min (examples at different time lags are shown in **Extended data Fig. 7**). The dots indicate data from a single-track segment. The dashed lines serve as visual guides. Data were pooled from three independent experiments for imDC and mDC.

Next, we quantitatively analyzed the migration characteristics of DCs in the three modes (**Supplementary Section F**). The turning angle heat map shows the change in directional persistence with time lag Δ*t* (**Fig. 3c, d**). Up to Δ*t* ∼ 3 min, the turning angle *θ* showed a peak at 0, indicating that the cells keep moving in the same direction. This was true for all three modes. In contrast, for Δ*t* > 3 min, the SD migration mode exhibited a strong anti-persistent memory with two peaks at *θ* ≈ ±*π*, which extended over several tens of minutes, resulting in subdiffusive migration with an anomalous exponent *α* = 0.6 (**Fig. 3e, Extended data Fig. 4**). SD migration showed a non-Gaussian diffusion pattern (**Extended data Fig. 5**) and differs from a zigzag-type anti-persistent walk. As shown in the phase-space density plot for the turning angle difference (Δ*θ*_*n*_) and radial velocities (*V*_*n*_) (**Fig. 3f, Extended data Fig. 6, 7**), *P*(Δ*θ*_*n*_) shows a peak at 0 and *P*(*V*_*n*_) is symmetric at 0, indicating that successive displacements preserve the same turning angles. Accordingly, SD migration was curly, as displayed in the exemplified trajectories (**Fig. 3b**).

The SP mode exhibited a zigzag-like persistent walking pattern (**Fig. 3b**). Interestingly, in the turning angle heat map, DCs show features of both persistent (*θ* ≈ 0) and anti-persistent (*θ* ≈ ±*π*) walks (**Fig. 3d**). This seemingly contradictory feature was obtained from single-cell motion analysis, and is not a superposition of two distinct cell motions (**Extended data Fig. 6**). The mean-squared displacement (MSD) analysis shows that SP migration is superdiffusive with *α* ≈ 1.5 for Δ*t* >10 min (**Fig. 3e**). For Δ*t* < 10 min, the SP mode of migration was similar to that in the SD mode. SP cell migration was highly heterogeneous (**Extended data Fig. 4**), and showed non-Gaussian diffusion with a displacement probability density function (PDF) *P*(Δ*x*|*t*), similar to that of SD motion (**Extended data Fig. 5**). The asymmetric *P*(*V*_*n*_) showed a positive part larger than the negative part indicating a persistent sequential walk (**Fig. 3f**). Further analyses using the phase-space maps and zigzag preference factor demonstrated that SP migration shows an empirically observed zigzag pattern (**Extended data Fig. 7-9**).

The FP migration mode showed a strong directional walk (**Fig. 3b**). The turning angle plot suggests that migration directionality is maintained for the total length of the trajectory (60 min) without significant dispersion of the angle fluctuation ⟨*θ*^2^⟩ with increasing time (**Fig. 3d**). FP migration is not ballistic but superdiffusive with *α* ≈ 1.7 (**Fig. 3e**), indicating faster and more directional motility than the SP mode. Unlike the SD and SP modes, FP migration dynamics are homogeneous (**Extended data Fig. 4**). Interestingly, *P*(Δ*x*|*t*) is a Laplace distribution, which indicates that a power-law tail does not exist. Actively diffusing biological particles often exhibit a power-law PDF of displacement as a signature of Lévy walks. Examples include *Escherichia coli* (i.e., run-and-tumble micro-swimmers)^5^, motor-driven macromolecules in the cytoplasm^24^, mRNA-protein complexes^25^, foraging birds^26^, and chemokine-stimulated immune cells (CD8+ T cell)^6^. However, the results of our analysis indicate that DC migration dynamics do not belong to the class of Lévy walks, regardless of maturation (**Extended data Fig. 5**).

To investigate how the maturation status of DCs influences the distribution of the three migration modes, we assigned the migration mode 1 h interval traces over time until the cell finally escaped from the field of view (**Fig. 4a, Supplementary Videos 6,7**). The speed and fraction of the three migration modes suggest that DC migration occurs mostly in the SD and SP modes, and their speeds are unaffected, regardless of their maturation status (**Fig. 4b,c**). The FP mode was more frequently observed in mDCs than in imDCs.

**Fig. 4.**
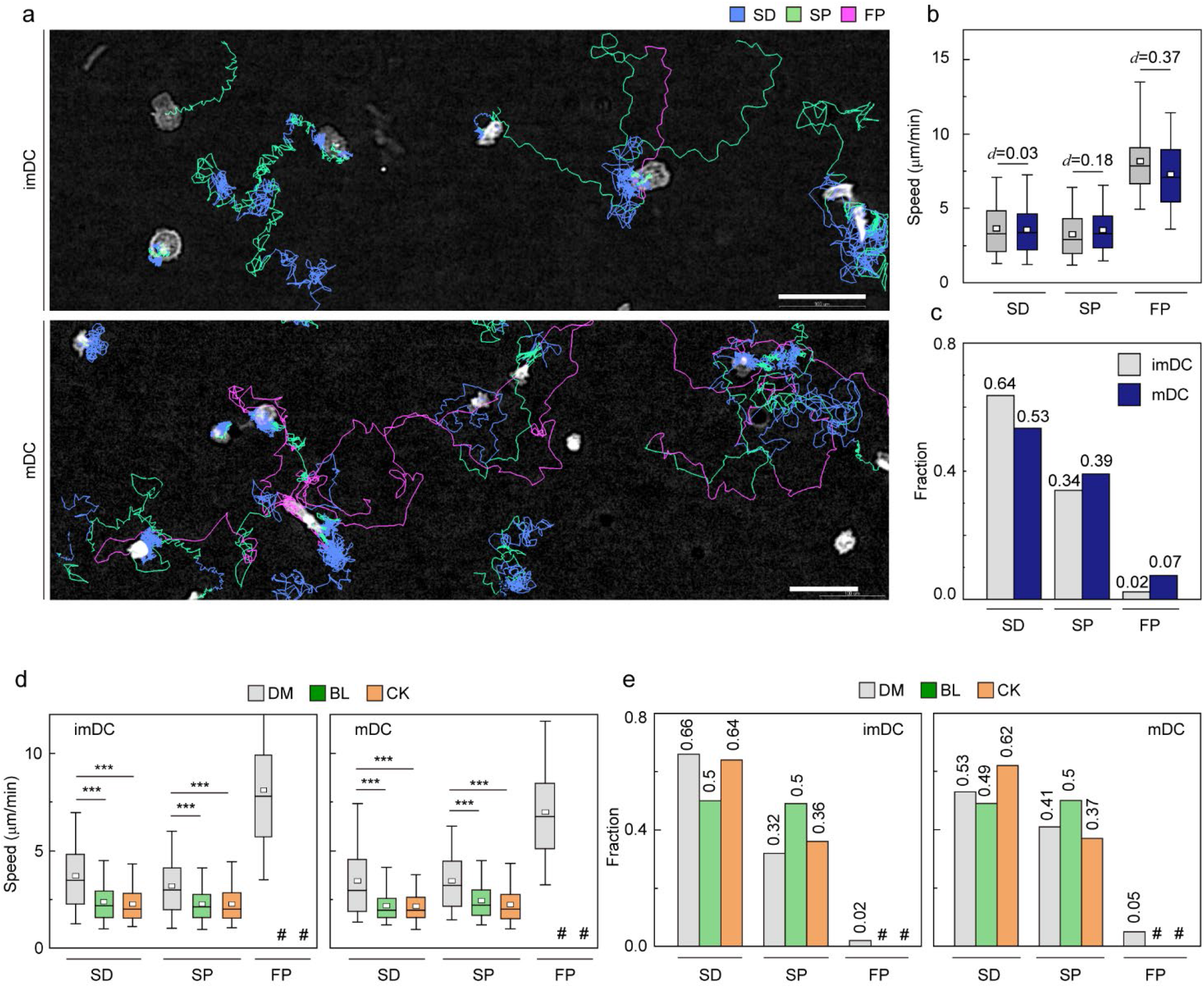
Distribution of migration modes. **a**, Representative images depicting the mode assignment for the individual cell migration track. Each line represents the migration mode for the cellular migration trajectories, with colors indicating SD (blue), SP (green), and FP (red) modes. Background-subtraction was performed to increase the contrast of brightfield image (scale bar = 100 μm). **b**, Mode mean track speed of imDCs and mDCs. Data were pooled from three independent experiments. (SD: imDC n=3660; mDC n= 1624, SP: imDC n=1953; mDC n=1192, FP: imDC n=132; mDC n=226). Cohen’s d (d) was used to indicate the standardized difference between the two means. **c**, Occurrence frequency of SD, SP, and FP modes in the imDC and mDC motility data. **d**, Mean track speed of imDCs and mDCs after treatment with DMSO (0.1%), 20 μM blebbistatin (BL, myosin II inhibition), or 100 μM CK666 (CK, Arp2/3 inhibition). In the box plots, the bars include 95% of the points, the center corresponds to the median, and the box contains 75% of the data. Data were pooled from three independent experiments. (imDC: SD, DM n=3702, BL n=1491, CK n=2910; SP, DM n=1815, BL n=1508, CK n=1613; FP, DM n=99, BL n=11, CK n=15; mDC: SD, DM n=1588, BL n=607, CK n=973; SP, DM n=1228, BL n=612, CK n=584; FP, DM n=162, BL n=10, CK n=9) The Kruskal-Wallis test with Dunn’s post hoc analysis was used to determine statistical significance. ***: *P* <0.001. #: very low incidence. **e**, Occurrence frequency of SD, SP, and FP modes for imDC and mDC migration after drug treatment.

Cell speed and persistence are controlled by actin polymerization dynamics^27,28^. Because myosin II-mediated contractility and Arp2/3-related actin polymerization are essential mediators of cell motility^29^, we sought to test how these molecules influence migration mode distribution (**Supplementary Videos 8–11**). Although the overall speed of both imDCs and mDCs was reduced, which is consistent with previous studies (**Extended data Fig. 10b**)^14,30–32^, the FP mode was absent, and the mean speed of both SD and SP modes was decreased significantly for both cases of myosin II and Arp2/3 inhibition (**Fig. 4d**). Remarkably, myosin II inhibition increased the SP mode of migration in the imDC population (**Fig. 4e**), whereas Arp2/3 inhibition increased the SD mode of migration of mDCs but had a negligible effect on imDCs. Thus, our findings provide microscopic explanation on the recent studies showed that inhibiting myosin II resulted in increased persistent migration^14,31,33^. The minimum change in the mode fractions of mDCs may result because they show sustained persistent motility^12,13^. Unstable lamellipodium formation caused by Arp2/3 inhibition impedes motility persistence^14,34^. Because mDCs have a relatively high SP mode fraction, inhibition of persistent motility appears to be more striking. Furthermore, myosin II inhibition decreased the imDC lifetime in the SD mode but increased the SD and SP mode lifetimes of mDCs. Arp2/3 complex inhibition increased the lifetime of mDCs in the SD mode, but not that of imDCs, indicating that mode lifetime dynamics and the role of actin polymerization are highly dependent on the maturation status of DCs (**Extended data Fig. 10**).

To study the transition dynamics of the DC migration modes, we constructed a transition matrix (**Fig. 5a**) that showed several noteworthy features. First, self-transition predominates, signifying that DCs move with strong mode persistence. Compared with imDCs, mDCs have higher self-transition rates for the SP and FP migration modes, and lower self-transition rates for the SD migration mode. This tendency explains why mDCs moved more persistently than imDCs; mDCs avoided SD migration in favor of SP or FP migration. Second, SD was the most recurrent migration mode for both imDCs and mDCs. The highest sum of influx rates was observed for the SD mode (∑_*i*∈{SD,SP,FP}_*P*_*i*→*SD*_). Third, maturation increases the transition to FP mode. Comparing the FP column of the transition matrix of imDCs and mDCs, we found that every component of the transition from SD, SP, and FP to FP was significantly increased.

**Fig. 5.**
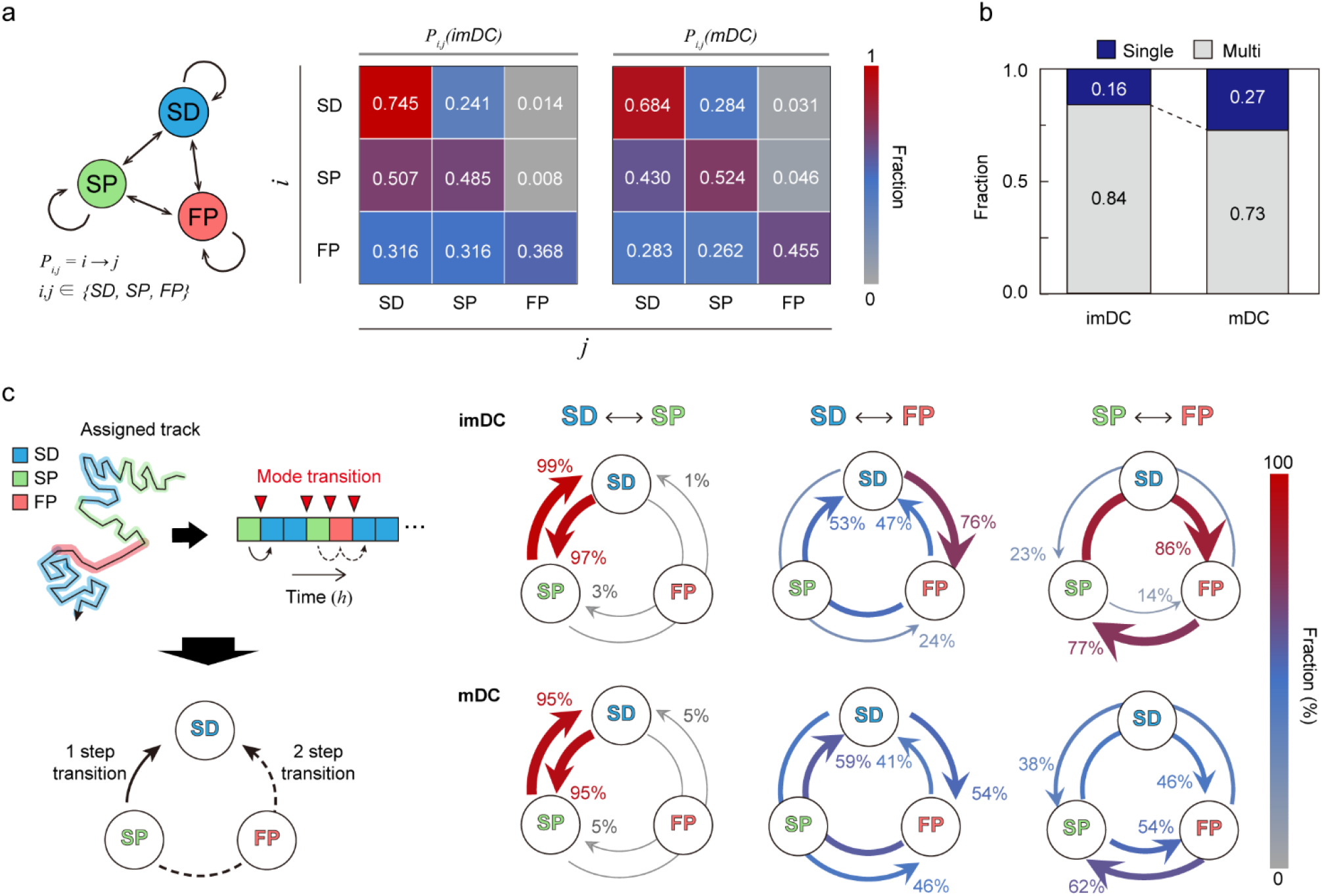
Transition dynamics of migration modes. **a**, Transition model between the three migration modes. *P*_*ij*_ denotes the transition rate from modes *i* to *j* (including self-transition) where *i, j* ∈ {*SD, SP, FP*}. **b**, Fraction of homogeneously (non-transitionary) and heterogeneously (transitionary) migrating cells over the observed duration. **c**, The time trace of migrating modes is depicted graphically, as well as the time points at which the mode shift occurs (red inverse triangle). The mode transition may occur in one step (e.g., SP→SD) and two steps (e.g., SP→FP→SD). The probabilities of a one-step or two-step mode transition from modes *i* to *j* (*i* ≠ *j*) were evaluated and plotted together for the cross-transition between different migration modes. The transition, including the FP mode, shows distinct pathways depending on maturation level. imDCs showed the cyclic mode transition (SD→FP→SP→SD), whereas mDCs had a more bidirectional transition. Data were pooled from three independent experiments. All plots shown here were obtained from three independent experiments.

Irrespective of maturation, DC migration is dominated by two slow modes (SD and SP), which suggests that DCs usually move at a slow speed with or without directional persistence. The high energy-consuming FP migration mode was observed intermittently, and the frequency of occurrence increased after maturation. In **Fig. 5b**, we plotted the population fraction of mode-preserving (single-mode) and mode-changing (multi-mode) cellular migration tracks. Interestingly, mDCs have a larger fraction of single-mode migration tracks than imDCs, indicating that maturation is involved in the stabilization of the innate migration of DCs.

Next, we examined the cross-transition dynamics among different migration modes (**Fig. 5c**). Mode changes can occur in one or two steps. We emphasize that the cross-mode transition diagram should be distinguished from the total transition matrix, as shown in **Fig. 5a**. For both imDCs and mDCs, a single-step change was the dominant pathway for the transition between the SD and SP modes. For the transition from the SD to FP modes, the one-step pathway was the most common for imDCs. For mDCs, the fractions for the one-step (SD→FP) and two-step (SD→SP→FP) transitions were similar. From the SP to FP transition, the two-step transition (SP→SD→FP) was the primary pathway for imDCs, whereas both one- and two-step transitions were similar for mDCs. Taken together, imDCs predominantly followed the unicyclic SD→FP→SP→SD transition (**Supplementary Video 12**). In contrast, mDCs showed no transition directionality. Based on these findings, we hypothesized that imDCs may evacuate the antigen-cleared location quickly and subsequently slow down gradually to find a new site to settle-down before displaying diffusive motility to acquire antigens. Even though the time scale was much shorter (< 30 min), faster acceleration than deceleration of imDC motility was noted in the 1D channel^35^. Although persistence-speed coupled biphasic intermittent random walks have been shown to boost the search efficiency of imDCs^36^, further analysis of the roles of the SP mode in the deceleration process is required.

Overall, our results suggest that ML-assisted analysis can successfully identify the heterogeneous motility patterns of DCs. Our findings indicate that the overall diffusive and persistent cellular motion of DCs is an emergent property of the dynamic switching of the three migratory modes, SD, SP, and FP, and are controlled by intracellular actin dynamics. Furthermore, the history-dependent mode transition of motile cells may offer a new paradigm for understanding complex cellular motility as an alternative to the current analysis of memoryless diffusive particles. This approach can be further utilized to understand the complex dynamics of cellular migration in response to external stimuli, such as environmental stiffness, physical confinement, and pathogens.

## Acknowledgments

This study was supported by taxpayers of South Korea through the Institute for Basic Science (project code IBS-R020-D1). T.S. and J.-H.J. acknowledge the financial support by the National Research Foundation of Korea (2022R1F1A1074045, 2017K1A1A2013241, 2020R1A2C4002490). We appreciate the manual trajectory correction of Ms. Yeojin Lee (UNIST, Ulsan, Korea) and Ms. Jae-Eun Kwon (UNIST, Ulsan, Korea).

## Author contributions

All authors designed the research, analyzed data, and wrote the paper. Y.C. performed the experiments and analyzed the data. T.S. performed the machine learning analysis with input by all authors. J.-H.J. and Y.-K.C. supervised the project.

## Additional information

### Extended data

Extended data is available for this paper at

### Supplementary information

Supporting information is available for this paper at

### Reprints and permissions information

Available at

### Data availability

The data and analysis code that support the findings of this study are available at:

---

**Extended Data Fig. 1.**
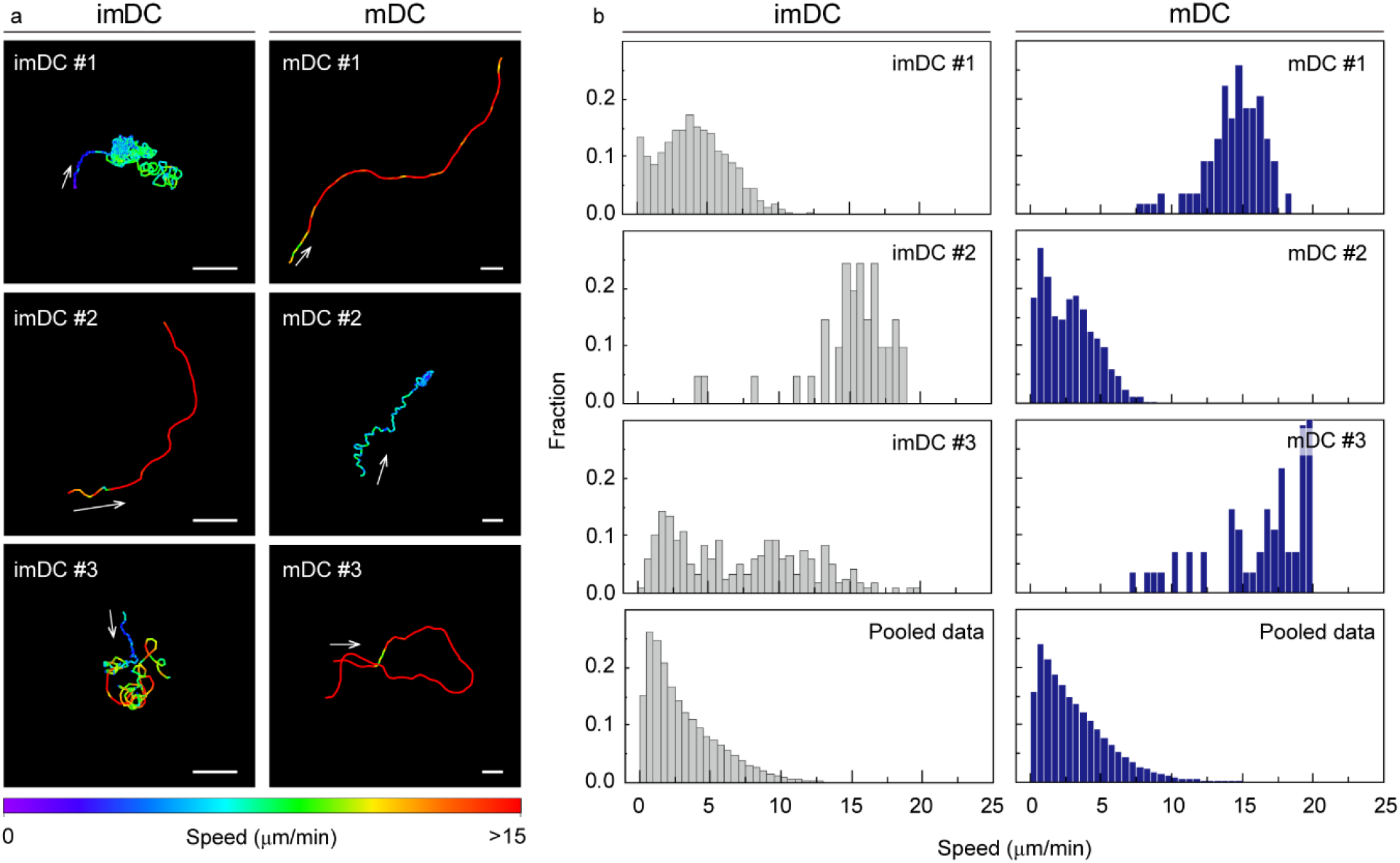
DC heterogeneous tracks and missing minor tracks in the averaged result. Even in the same maturation status, atypical motilities are exhibited in the population. **a**, Additional representative tracks are plotted with color codes. Color codes indicate the instantaneous speed. **b**, Distribution of instantaneous speed from **a**. The single-cell scale tracks showed heterogeneous motility patterns; however, this heterogeneity was absent in the averaged results from whole cells (pooled data). Single-cell tracks were collected from one of three experiments (imDC: n=93, mDC: n=94).

**Extended Data Fig. 2.**
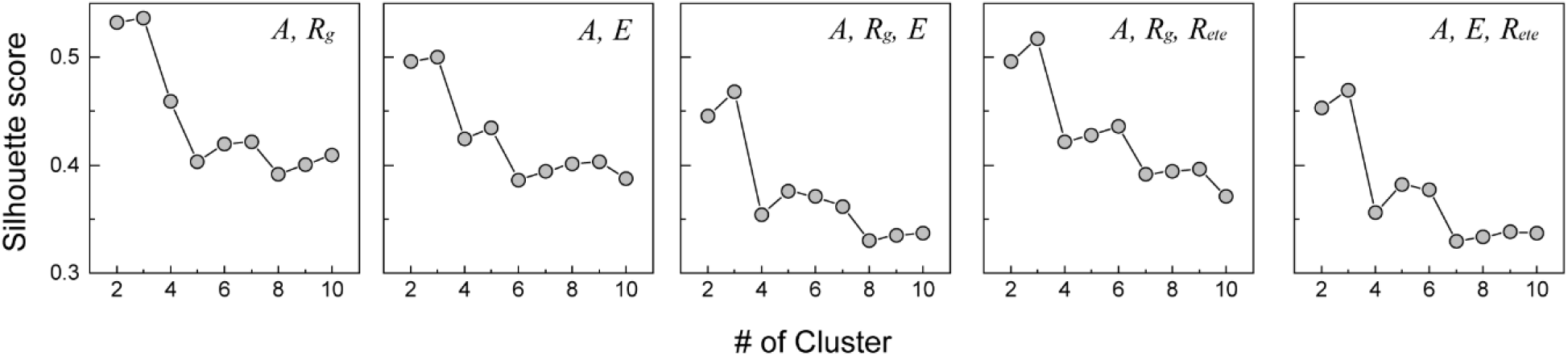
The number of distinct groups obtained using an averaged silhouette score. The number of groups is shown as a function of the number of clusters using K-means clustering with five distinct combinations of the original features. The feature combination is annotated in the legend. In the plots, all features were normalized using the MinMaxScaler in the range of [**0, 1**] provided by the scikit-learn package in Python^23^.

**Extended Data Fig. 3.**
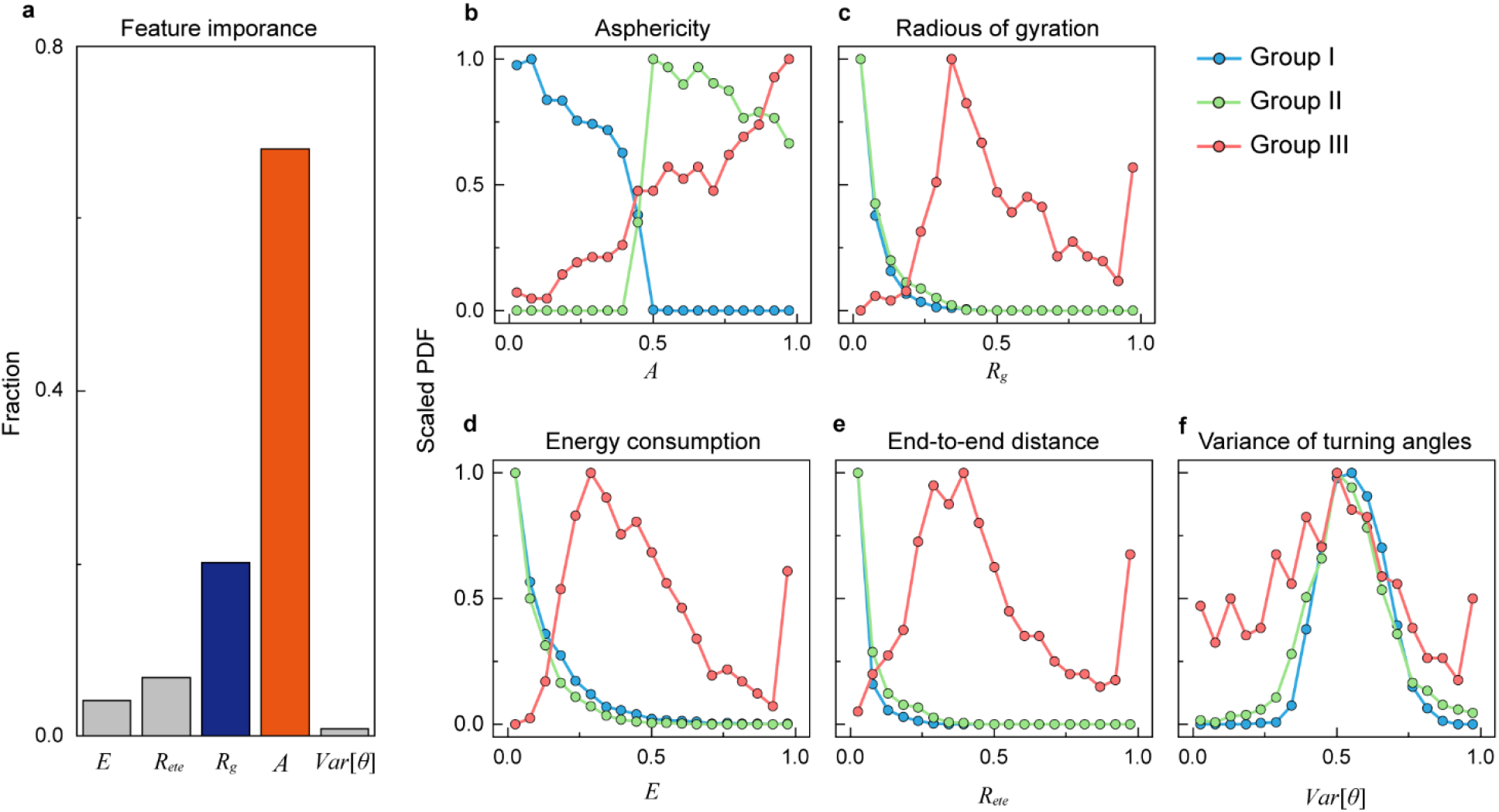
Feature importance and feature distribution for three groups. **a**, The feature importance (%) relative to the five features used in machine learning kernel training. The trained algorithm extracts the distributions of five features for each group, including Asphericity (**b**), Radius of gyration (**c**), End-to-end distance (**d**), Energy consumption (**e**), and Variance of turning angles (**f**). The color indices are blue (group I), green (group II), and red (group III).

**Extended Data Fig. 4.**
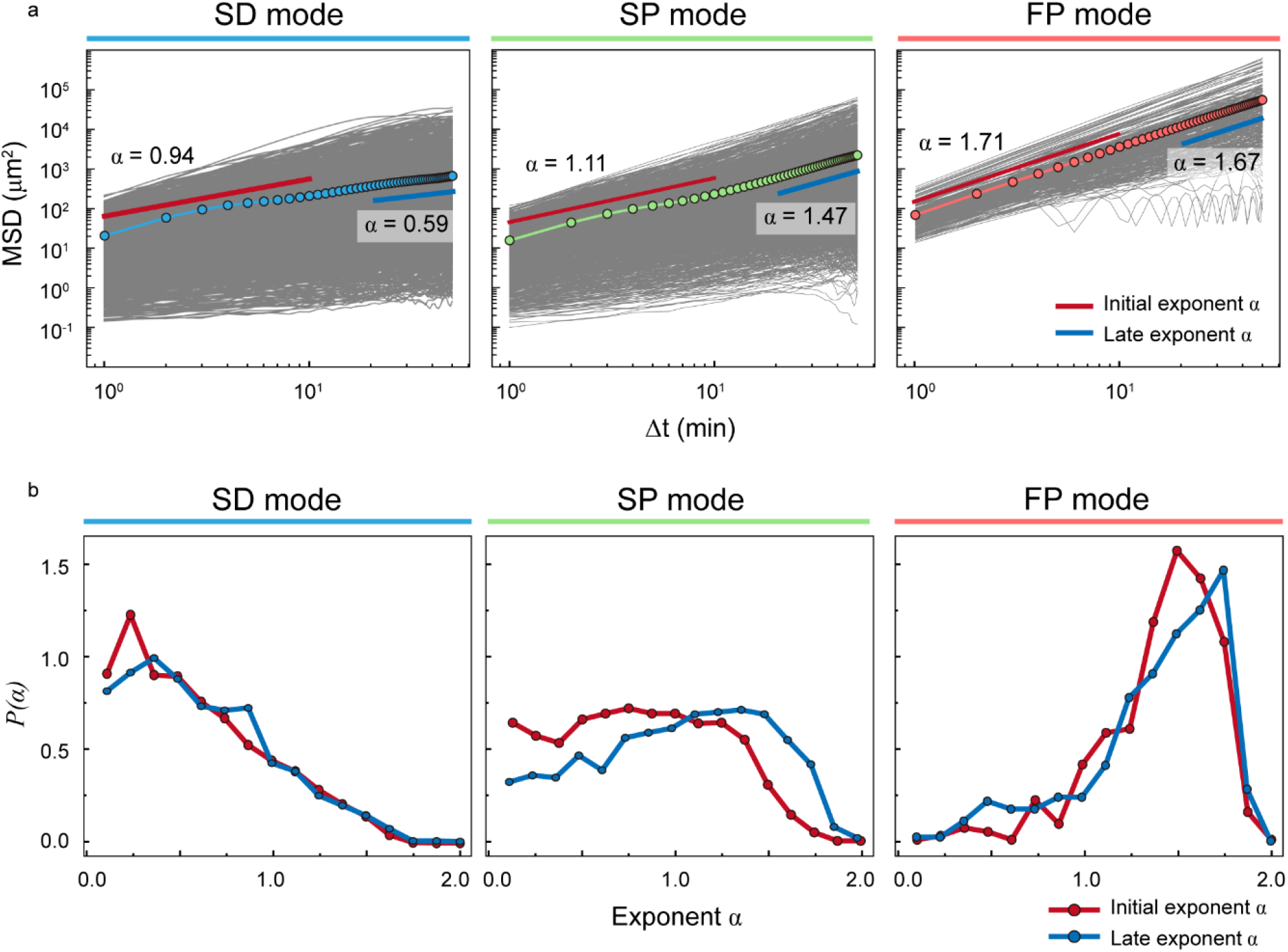
The MSD curves and the distribution of anomalous exponent *α* for the SD, SP, and FP modes. **a**, In the MSD plot, the gray lines depict the individual MSD curves and the symbol with the solid line indicates their average. We obtained two fitted anomalous exponents from the averaged MSD: The short-time exponent for Δ*t* ∈ [0,10] min (initial) and the long-time exponent for Δ*t* ∈ [20,50] min (late). **b**, The distribution of the fitted anomalous exponents ***α*** from the individual MSD curves in **a**. Each panel shows two *P*(*α*) for the short- and long-time regimes.

**Extended Data Fig. 5.**
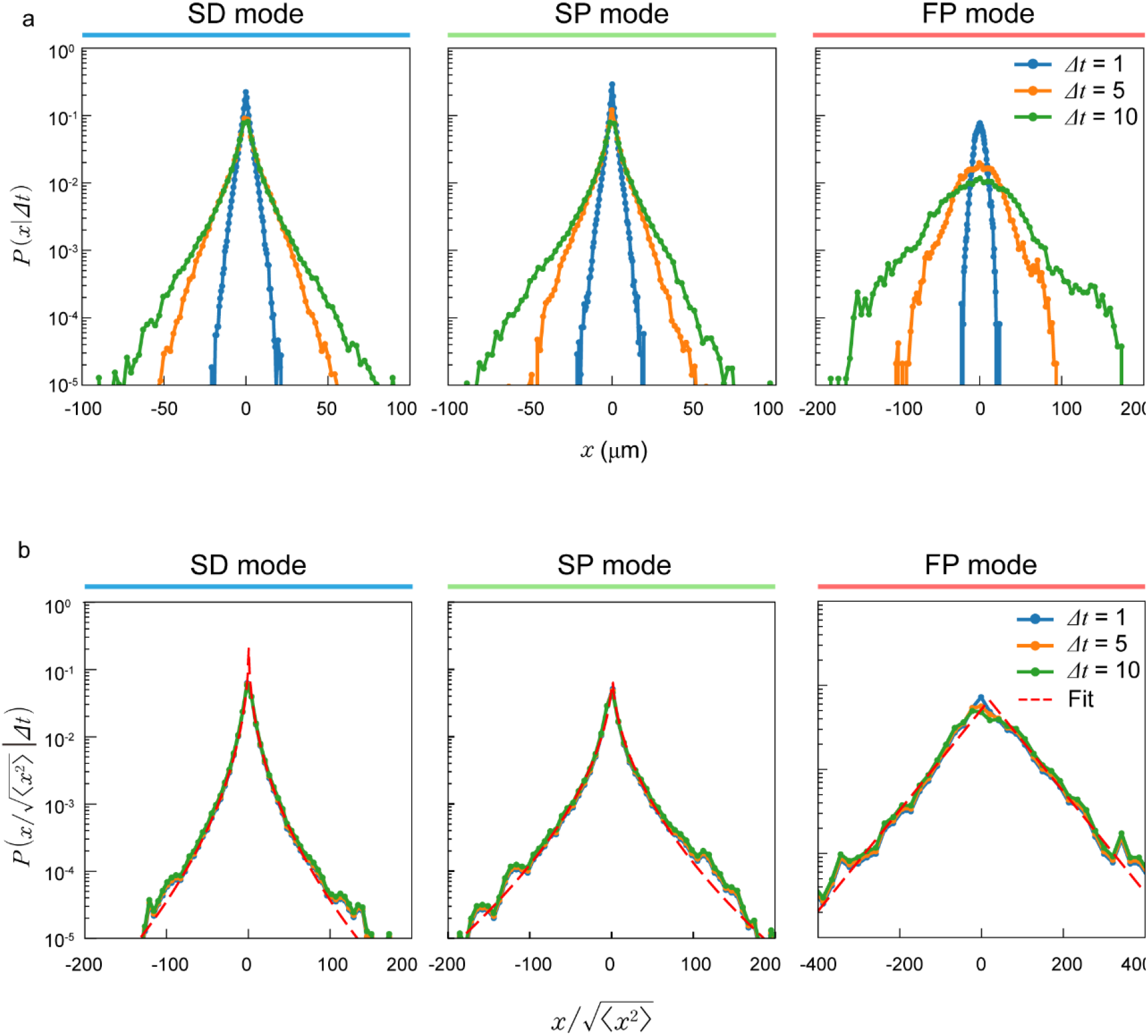
Displacement probability density functions (PDFs) *P*(*x*|Δ*t*) and their re-scaled PDFs. **a**, The displayed PDFs show the *x*-component displacement. Similar results were obtained for *y*-component displacements. The three colors represent the lag time indicated in the legend of the last column. **b**, The re-scaled PDFs were obtained by plotting the original PDFs with 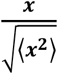. We fitted the re-scaled PDF with a stretched-exponential function 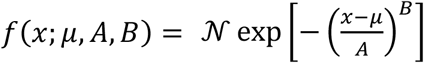. The red dashed line indicates the best fit with the parameters (*μ, A, B*) : (0.48,1.07,0.48) (SD), (1.09,2.84,0.53) (SP), and (18.23,75.26,0.98) (FP), respectively. The stretched exponent *B* is less than unity, indicating that DC migration is highly deviant from Gaussian dynamics (*B* = 2).

**Extended Data Fig. 6.**
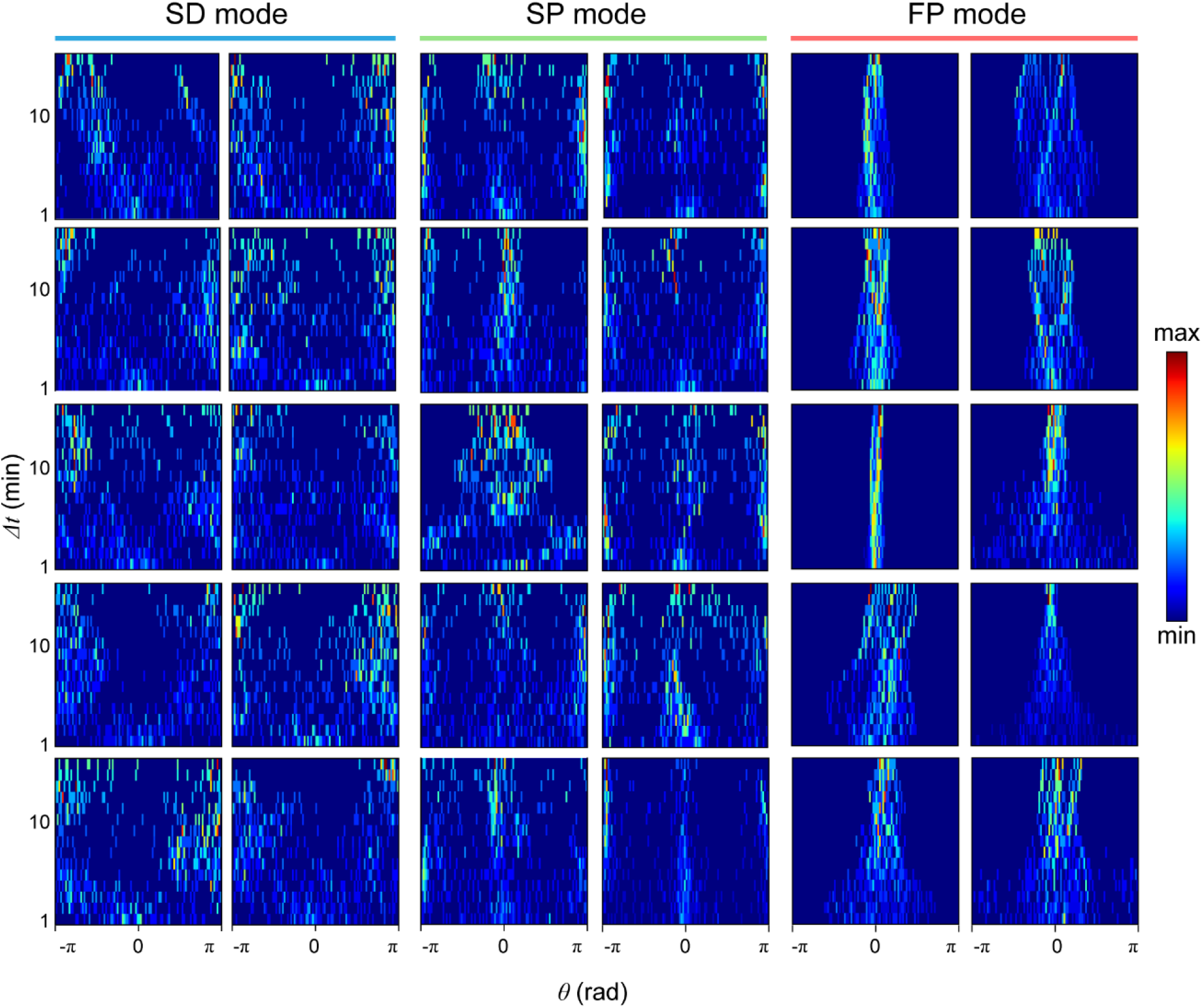
Turning angle heat maps from randomly selected individual trajectories. For each mode, ten samples were analyzed.

**Extended Data Fig. 7.**
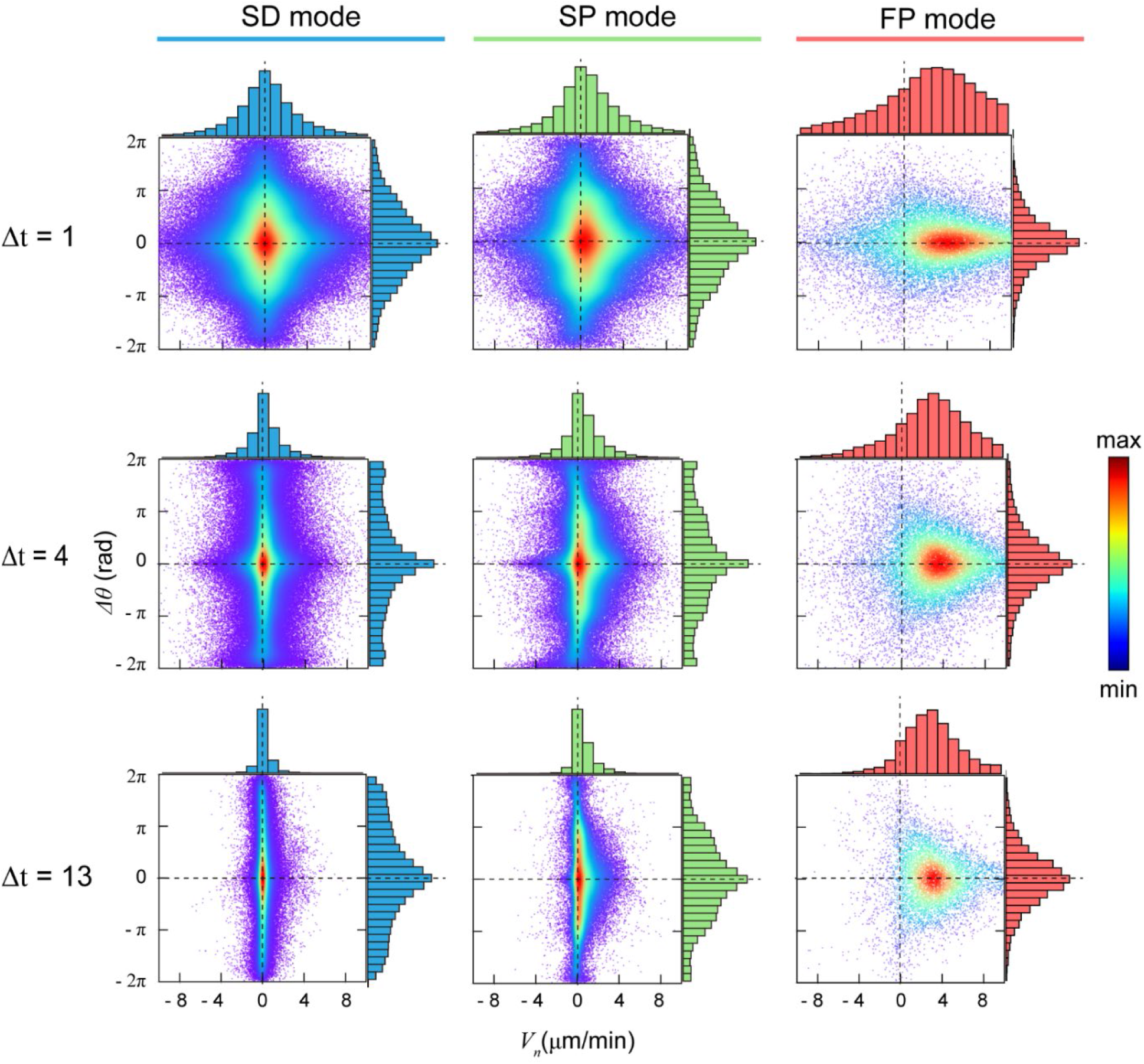
The density map of the phase-space space (Δ*θ*_*n*_, *V*_*n*_)for the three migration modes. For each mode (column), color-coded density maps at three lag times (row, min) were plotted. The dots indicate data from a single-track segment. The dashed lines serve as visual guides.

**Extended Data Fig. 8.**
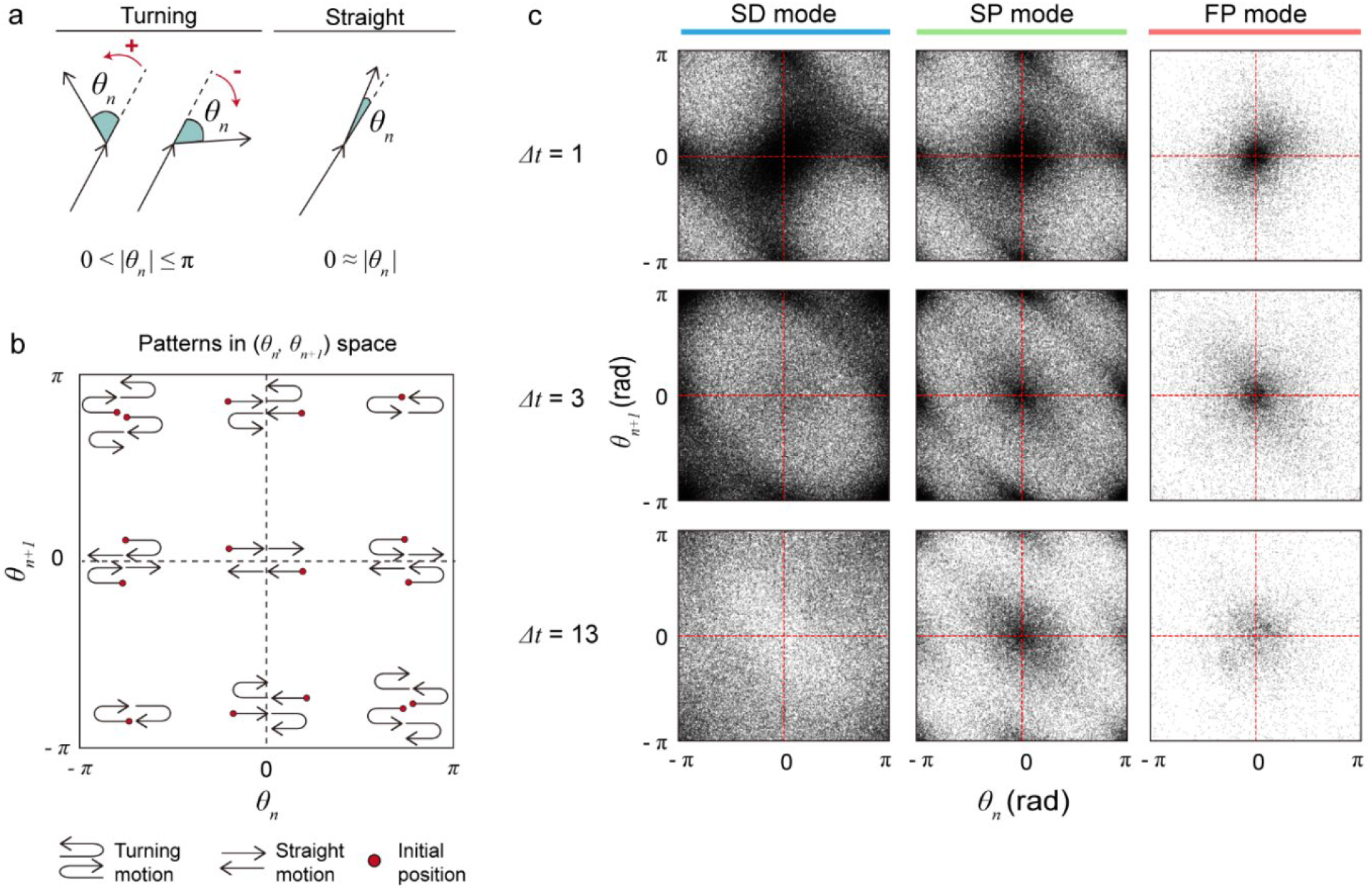
Density maps of the phase-space (*θ*_*n*_, *θ*_*n*+1_)for various lag times. **a**, Cartoon showing turning and straight movement. The possible range of ***θ***_***n***_ is shown. **b**, The nine dense spots in **c** and the corresponding trajectory motif. **c**, The scatter points are collected from all trajectory samples for a given mode over the entire period. The dots indicate data from a single-track segment. The dashed lines serve as visual guides.

**Extended Data Fig. 9.**
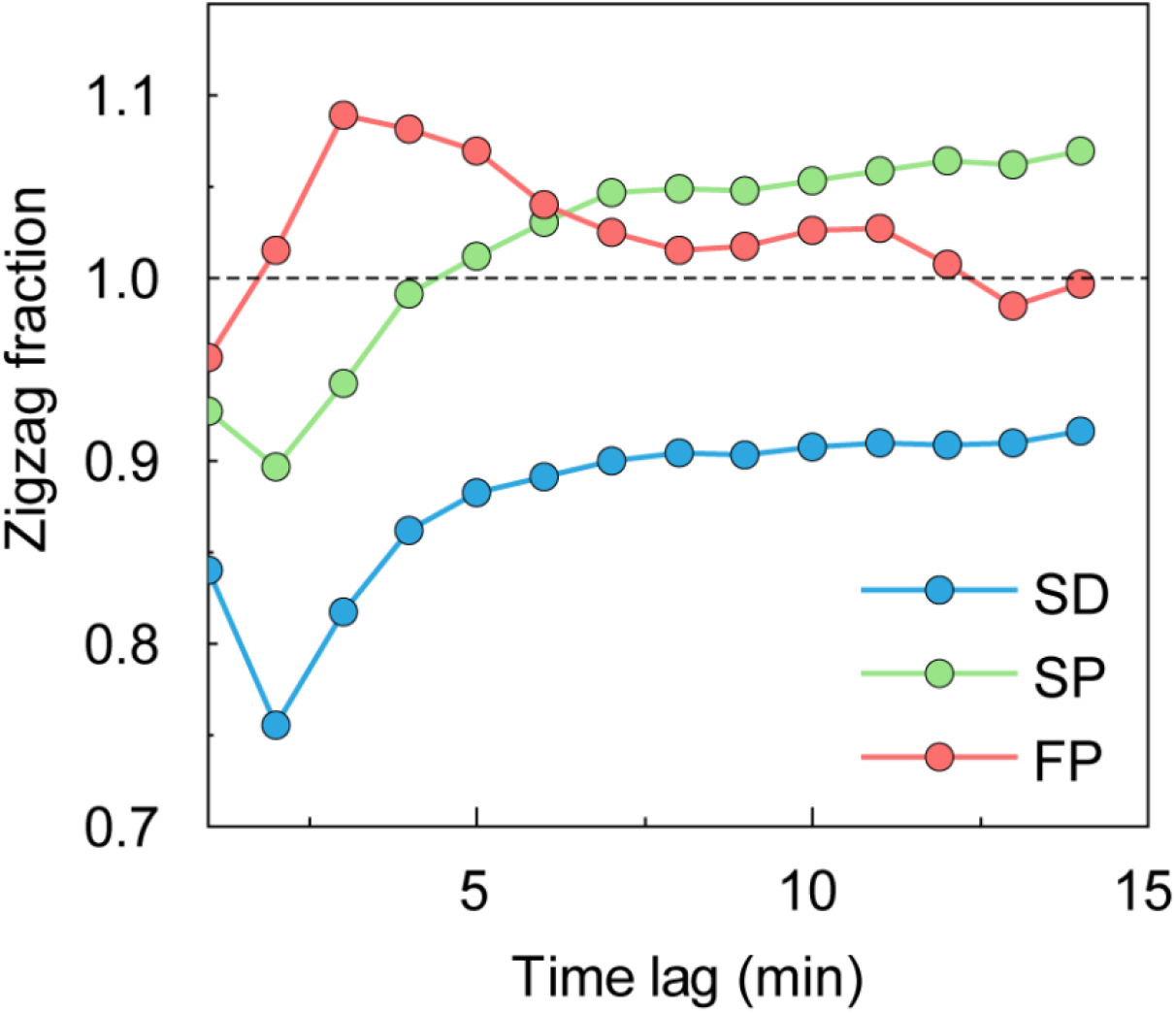
The zigzag fraction as a function of the lag time. The zigzag fraction is the ratio of the total number of scatter points between the first-third quadrant and the second-fourth quadrant in the density map. The color code represents the SD (blue), SP (green), and FP (red) modes. When the fraction becomes unity (the dashed line), it signifies that the cell has the same amount of two sequential curved events over the entire migration: zigzag-like opposite turning events (left-right or right-left turn) and circular motion (left-left or right-right turn).

**Extended Data Fig. 10.**
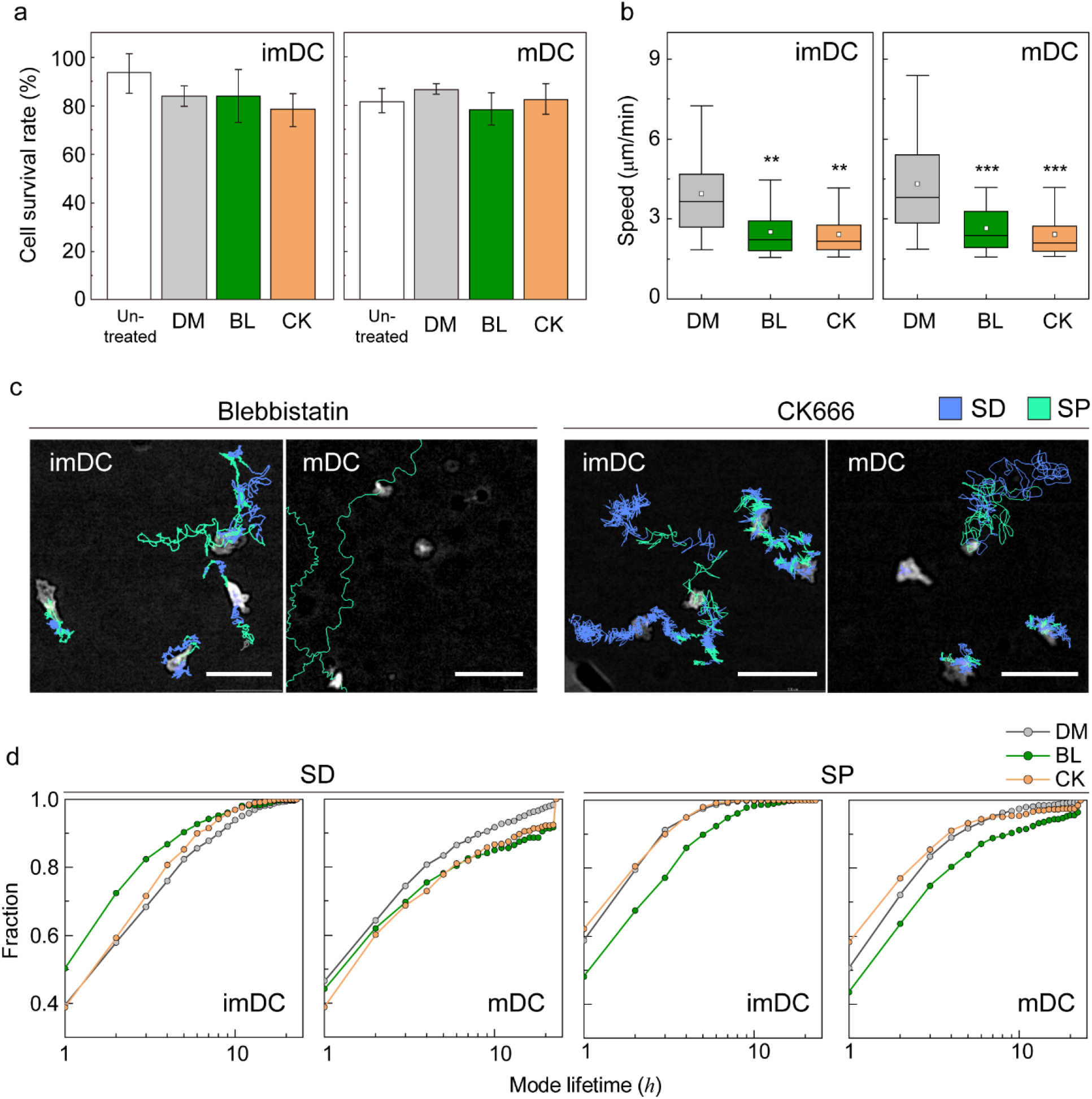
Molecular inhibition in DC migration experiments. **a**, Cell survival rate of imDCs and mDCs at 24 h after drug treatments. DM: DMSO (0.1%), BL: blebbistatin (20 μM), and CK: CK666 (100 μM). **b**, Mean track speed of imDCs and mDCs with drug treatments. In the box plots, the bars include 95% of the points, the center corresponds to the median, and the box contains 75% of the data. Data were pooled from three independent experiments. The Kruskal-Wallis test with Dunn’s post hoc test was used to determine statistical significance. **: *P* < 0.01; ***: *P* < 0.001. (imDC: DM n=444, BL n=213, CK n=286, mDC: DM n=321, BL n=102, CK n=121) **c**, Representative images of imDCs and mDCs under the effect of drugs. Each line represents the migration mode for the cellular migration trajectories with colors indicating SD (blue), and SP (green) modes. Background-subtraction was performed to increase the contrast of brightfield image (scale bar = 100 μm). **d**, Cumulative distribution of consecutive mode lifetime for imDC and mDC migration after drug treatment.

## References

1. Vicente-Manzanares, M., Webb, D. J. & Horwitz, A. R. Cell migration at a glance. J Cell Sci 118, 4917–9 (2005).

2. Dieterich, P., Klages, R., Preuss, R. & Schwab, A. Anomalous dynamics of cell migration. Proc Natl Acad Sci U A 105, 459–63 (2008).

3. Bechinger, C. et al. Active Particles in Complex and Crowded Environments. Rev. Mod. Phys. 88, (2016).

4. Metzler, R. & Klafter, J. The random walk’s guide to anomalous diffusion: a fractional dynamics approach. Phys. Rep. 339, 1–77 (2000).

5. Ariel, G. et al. Swarming bacteria migrate by Levy Walk. Nat Commun 6, 8396 (2015).

6. Harris, T. H. et al. Generalized Levy walks and the role of chemokines in migration of effector CD8+ T cells. Nature 486, 545–8 (2012).

7. Moreau, H. D., Piel, M., Voituriez, R. & Lennon-Dumnil, A.-M. Integrating Physical and Molecular Insights on Immune Cell Migration. Trends Immunol. 39, 632--643 (2018).

8. Worbs, T., Hammerschmidt, S. I. & Forster, R. Dendritic cell migration in health and disease. Nat Rev Immunol 17, 30–48 (2017).

9. Chabaud, M. et al. Cell migration and antigen capture are antagonistic processes coupled by myosin II in dendritic cells. Nat Commun 6, 7526 (2015).

10. Faure-André, G. et al. Regulation of Dendritic Cell Migration by CD74, the MHC Class II-Associated Invariant Chain. Science 322, 1705–1710 (2008).

11. Solanes, P. et al. Space exploration by dendritic cells requires maintenance of myosin II activity by IP3 receptor 1. EMBO J 34, 798–810 (2015).

12. Vargas, P. et al. Innate control of actin nucleation determines two distinct migration behaviours in dendritic cells. Nat Cell Biol 18, 43–53 (2016).

13. Choi, Y., Sunkara, V., Lee, Y. & Cho, Y. K. Exhausted mature dendritic cells exhibit a slower and less persistent random motility but retain chemotaxis against CCL19. Lab Chip 22, 377–386 (2022).

14. Stankevicins, L. et al. Deterministic actin waves as generators of cell polarization cues. Proc Natl Acad Sci U A 117, 826–835 (2020).

15. Li, H. et al. Zigzag Generalized Levy Walk: the In Vivo Search Strategy of Immunocytes. Theranostics 5, 1275–90 (2015).

16. Choi, Y., Kwon, J. E. & Cho, Y. K. Dendritic Cell Migration Is Tuned by Mechanical Stiffness of the Confining Space. Cells 10, 3362 (2021).

17. Manzo, C. & Garcia-Parajo, M. F. A review of progress in single particle tracking: from methods to biophysical insights. Rep Prog Phys 78, 124601 (2015).

18. Newby, J. M., Schaefer, A. M., Lee, P. T., Forest, M. G. & Lai, S. K. Convolutional neural networks automate detection for tracking of submicron-scale particles in 2D and 3D. Proc Natl Acad Sci U A 115, 9026–9031 (2018).

19. Zhang, P. et al. Analyzing complex single-molecule emission patterns with deep learning. Nat Methods 15, 913–916 (2018).

20. Granik, N. et al. Single-Particle Diffusion Characterization by Deep Learning. Biophys J 117, 185–192 (2019).

21. Thapa, S., Lomholt, M. A., Krog, J., Cherstvy, A. G. & Metzler, R. Bayesian analysis of single-particle tracking data using the nested-sampling algorithm: maximum-likelihood model selection applied to stochastic-diffusivity data. Phys Chem Chem Phys 20, 29018–29037 (2018).

22. Lars Buitinck et al. API design for machine learning software: experiences from the scikit-learn project. ECML PKDD Workshop Lang. Data Min. Mach. Learn. 108–122 (2013).

23. Pedregosa, F. et al. Scikit-learn: Machine Learning in Python. J. Mach. Learn. Res. 12, 2825– 2830 (2011).

24. Chen, K., Wang, B. & Granick, S. Memoryless self-reinforcing directionality in endosomal active transport within living cells. Nat Mater 14, 589–93 (2015).

25. Song, M. S., Moon, H. C., Jeon, J. H. & Park, H. Y. Neuronal messenger ribonucleoprotein transport follows an aging Levy walk. Nat Commun 9, 344 (2018).

26. Viswanathan, G. M. et al. Levy flight search patterns of wandering albatrosses. Nature 381, 413–415 (1996).

27. Callan-Jones, A. C. & Voituriez, R. Actin flows in cell migration: from locomotion and polarity to trajectories. Curr Opin Cell Biol 38, 12–7 (2016).

28. Seetharaman, S. & Etienne-Manneville, S. Cytoskeletal Crosstalk in Cell Migration. Trends Cell Biol. 30, 720–735 (2020).

29. Kameritsch, P. & Renkawitz, J. Principles of Leukocyte Migration Strategies. Trends Cell Biol 30, 818–832 (2020).

30. Bretou, M. et al. Lysosome signaling controls the migration of dendritic cells. Sci Immunol 2, (2017).

31. Maiuri, P. et al. Actin flows mediate a universal coupling between cell speed and cell persistence. Cell 161, 374–86 (2015).

32. Gaertner, F. et al. WASp triggers mechanosensitive actin patches to facilitate immune cell migration in dense tissues. Dev Cell 57, 47–62 e9 (2022).

33. Allen, G. M. et al. Cell Mechanics at the Rear Act to Steer the Direction of Cell Migration. Cell Syst 11, 286–299 e4 (2020).

34. Krause, M. & Gautreau, A. Steering cell migration: lamellipodium dynamics and the regulation of directional persistence. Nat Rev Mol Cell Biol 15, 577–90 (2014).

35. Lavi, I., Piel, M., Lennon-Dumenil, A. M., Voituriez, R. & Gov, N. S. Deterministic patterns in cell motility. Nat. Phys. 12, 1146–1152 (2016).

36. Shaebani, M. R., Jose, R., Santen, L., Stankevicins, L. & Lautenschlager, F. Persistence-Speed Coupling Enhances the Search Efficiency of Migrating Immune Cells. Phys Rev Lett 125, 268102 (2020).

37. MacQueen, J. Some methods for classification and analysis of multivariate observations. Proc. Fifth Berkeley Symp. Math. Stat. Probab. Vol. 1 Stat. 5.1, 281–298 (1967).

38. Kanungo, T. et al. An efficient k-means clustering algorithm: analysis and implementation. IEEE Trans. Pattern Anal. Mach. Intell. 24, 881–892 (2002).

39. Chen, T. & Guestrin, C. XGBoost. (2016).

40. Borisov, V. et al. Deep Neural Networks and Tabular Data: A Survey. (2022) doi:10.48550/arXiv.2110.01889.

