## Supplementary Information for "Machine learning analysis reveals the dynamics of mode transition in dendritic cell migration"

[1] Department of Physics, Pohang University of Science and Technology (POSTECH), Pohang, 37673, Republic of Korea, [2] Department of Data information and Physics, Kongju National University, Gongju, 32588, Republic of Korea, [3] Center for Soft and Living Matter, Institute for Basic Science (IBS), Ulsan, 44919, Republic of Korea, [4] Biomedical Engineering, Ulsan National Institute of Science and Technology (UNIST), Ulsan, 44919, Republic of Korea, [5]Asia Pacific Center for Theoretical Physics (APCTP), Pohang, 37673, Republic of Korea

†These authors contributed equally to this work.
*

**This PDF file includes:**

Supplementary text
Figures S1 to S2
Legends for Video S1 to S12
SI References

**Other supplementary materials for this manuscript include the following:**

Videos S1 to S12

### A. Bone-Marrow derived Dendritic Cells (BMDCs)

#### 1) Cell culture

BMDCs were generated as described previously[1,2]. Briefly, tibias and femurs from BALB/c mice (8–12 weeks old, female) were flushed, and red blood cells were lysed with ammonium-chloride-potassium (ACK) lysis buffer (Gibco). Bone marrow cells were plated in 24 well tissue culture well plates (1 × $10^6$ cells/mL) in complete medium containing RPMI 1640, supplemented with 5% fetal bovine serum (FBS), 1% antibiotic-antimycotic solution, 1% HEPES buffer, and 0.1% 2-mercaptoethanol (all reagents were purchased from Gibco) containing 20 ng/mL recombinant mouse granulocyte-macrophage colony-stimulating factor (GM-CSF; Peprotech). The medium was completely replaced with fresh GM-CSF every two days. On day six, non-adherent and loosely adherent cells were collected by gentle pipetting and transferred to Petri dishes. After one day of culture, immature BMDCs, which appeared as floating cells, were collected. Phenol-red-free RPMI 1640 medium (Gibco) was used for fluorescence microscopy experiments, including cell height measurements and cell viability tests. To generate mature dendritic cells (mDCs), immature DCs (imDCs) were stimulated with 100 ng/mL lipopolysaccharide (LPS, LPS-EB Ultrapure; Invivogen) for 30 min. The cells were carefully washed three times and incubated for six hours in fresh complete medium, as described in previous studies[3]. After incubation, floating cells were harvested as mDCs. All animal experiments were performed according to protocols approved by the Institutional Animal Care and Use Committee of the Ulsan National Institute of Science and Technology (UNISTIACUC-19-15).

#### 2) DC characterization

The specific surface markers of imDCs and mDCs were characterized by flow cytometry (Cytoflex, Beckman Coulter) (**Fig. S1**). The upregulated expression levels of co-stimulatory molecules, CD86, CD80, and CD40, antigen-presenting molecule MHC class II (I-A/I-E), and the chemokine receptor CCR7 were evaluated as DC maturation markers, and CD11c and CD11b were analyzed as dendritic cell markers. The following antibodies were used in flow cytometry experiments: anti-CD86-FITC (GL1, 1:200),-CD40-PE (1C10, 1:200),-MHC class II (I-A/I-E)-FITC (M5/114.15.2, 1:200), and-CCR7-PE (4B12, 1:200) were purchased from Thermo Fisher Scientific, and anti-CD11b-APC (M1/70, 1:200), -CD11c-APC (HL3, 1:200), and-CD80-PE (16-10A1, 1:200) were purchased from BD Biosciences. The acquired data were analyzed using the FlowJo software (BD).

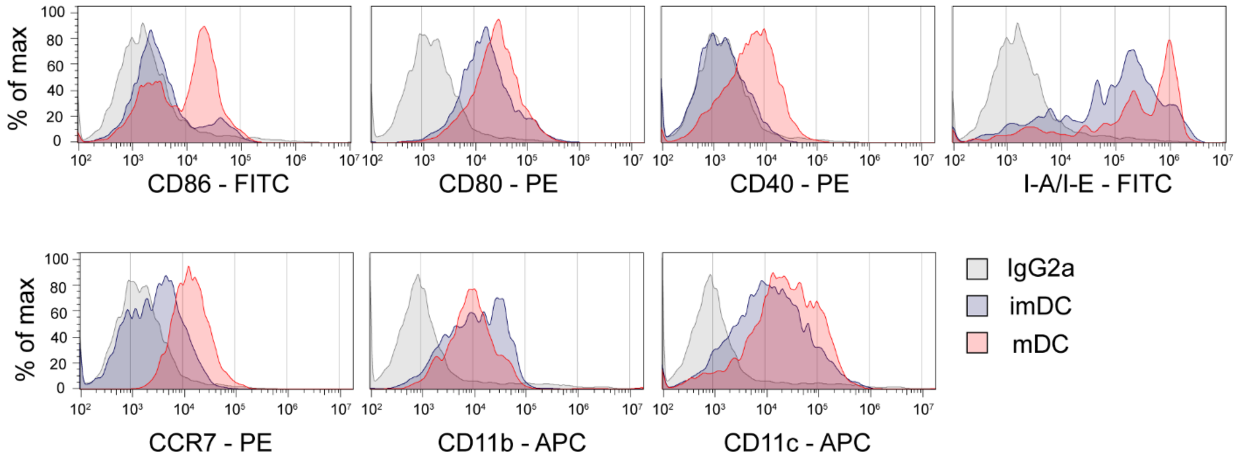


**Fig. S1. DC characterization.** Flow cytometry analysis was performed for all independent experiments to confirm the DC phenotype. Co-stimulatory molecules (CD86, CD80, and CD40), antigen-presenting molecules (MHC class II (I-A/I-E)), chemotaxis receptor (CCR7), and DC markers (CD11b and, CD11c) were evaluated in both imDCs and mDCs. One representative experiment out of three is shown.

#### B. Gel confiner

##### 1) Fabrication

Under-agarose migration by gel confinement was performed as described previously[1,2]. Briefly, the gel confiner consisted of a custom-designed PDMS structure and low-melting agarose gel. The PDMS structure consisted of a 10:1 ratio for the main body and a 30:1 ratio for the sticky PDMS-coated bottom. Before casting the gel solution, the PDMS structure was placed in a Petri dish. Low melting agarose (2.4%) was dissolved in phenol-red free HBSS buffer and heated at 80°C for 20 min, and the resulting solution was cooled at room temperature to 40°C. The same volume of 2x conditioned medium (RPMI 1640, 10% FBS, 2% HEPES buffer, 2% antibiotic-antimycotic solution, and 0.2% 2-mercaptoethanol) was mixed to a final concentration of 1.2% and cast to obtain the PDMS structure. The gel was cured for 20 min at room temperature. The cured gel confinement was incubated overnight in a cell culture incubator with complete medium. Subsequently, to prevent non-specific cell-to-cell interactions in the migration assay, 800 cells in a small drop of cell suspension were seeded on 10 mm diameter coverslips with 20 μg/mL bovine fibronectin-coating. The cells were incubated for 30 min in a cell culture incubator to enable them to settle on the substrate. Subsequently, the cells were carefully covered by gel confinement, and motility was imaged after 1 h.

##### 2) Measurement of Young's Moduli

Agarose gel blocks were prepared at concentrations of 1.2% (w/v), with the same gel confinement fabrication. The mechanical properties of the gels were determined using a rheometer (MCR502 WESP; Anton Paar). The gel height was approximately 1.0 mm. The shear moduli (G, Pa) were analyzed using the relationship between shear stress and shear strain before gel disruption, and Young's moduli (E, kPa) were calculated using the following equation: $E = 2(1 + v)G$, with the Poisson's ratio ($v$) of agarose set as 0.5[4].

##### 3) Measurement of cell height

To check the reproducibility of the confined environment, the height of the fluorescence-stained DCs under 2D gel confinement was measured using a laser scanning confocal microscope (A1R, Nikon) as shown in **Fig. S2**. The DC suspensions were stained using DiO (Invitrogen) for three minutes and washed thrice with complete medium. After three minutes of recovery, stained DCs were seeded onto the substrate and covered by gel confinement. The 3D confocal image was acquired using a 100x plan apo lens, and the *z* interval was 0.5 μm. Cell height was measured manually using an NIS Element (Nikon).

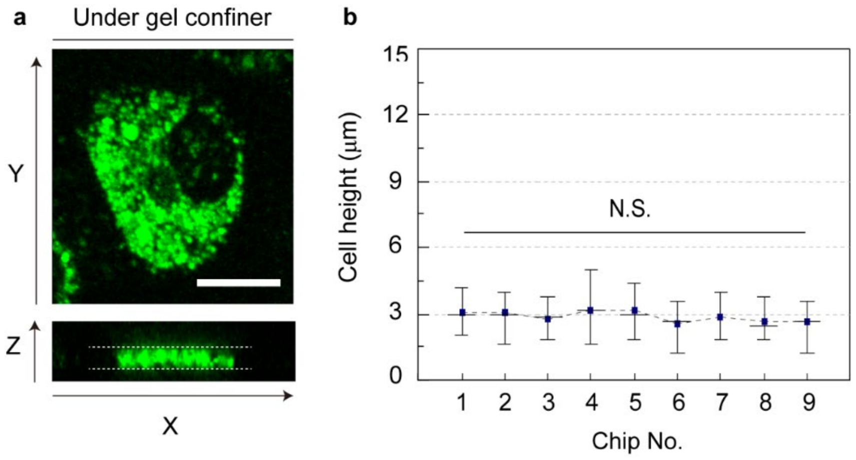

**Fig S2. Measurement of cell height. a.** 3D confocal microscopic image of DiO stained DCs. Cell height was measured to check the degree of confinement. **b.** Reproducibility of gel confiner. Nine independent chips were checked, and more than 20 cells at 5 different positions were investigated. In the plots, bars include 95% of the data, the central bars indicate the medians, and blue dots indicate the means. Kruskal–Wallis/Dunn's multiple comparisons testing was used to compare different sample populations; N.S.: $P > 0.05$.

#### C. Cell migration

##### 1) Live cell imaging

Live cell imaging was performed using an inverted microscope (Eclipse Ti-E, Nikon) configured with a 10x dry objective lens and sCMOS camera (Flash4.0, Hamamatsu). The cells were imaged for 24 h and bright-field images were obtained every 1 min. The recorded images were processed using the *z*-max intensity projection for non-labeled automatic tracking[5]. The sample focal plane was focused, and two more image sequences were obtained along the *z*-axis. Subsequently, rolling-ball background subtraction was performed, and the maximum intensity projection overlaid the *z*-stacks, facilitating contrast enhancement for cell segmentation. During the experiments, an incubator system (Chamlide HK; Live Cell Instrument) maintained the microscope at 37°C with 95% humidity and 5% $CO_2$.

##### 2) Cell migration tracking

For tracking, cells in the pre-processed image were automatically detected using the IMARIS (Bitplane) 'Spots' function. Cells were identified using intensity thresholding and size estimation as 20 μm diameter particles. The position of the center of mass was tracked, and tracking errors such as misconnections between different objects or misrecognized objects were corrected manually.

##### 3) Statistical analysis

All data represent the mean ± SEM of three independent experiments for each condition unless stated otherwise. Normality was determined using the D'Agostino and Pearson tests. The Mann-Whitney and Kruskal-Wallis tests along with Dunn's post hoc test were used to determine statistical significance. To prevent type I errors due to the large number of samples and confirm statistical significance, randomly selected samples were used to obtain the *P*-values. The criteria for random selection followed the hypothesized sample size, with a statistical power of 0.8. All statistical analyses were performed using the Origin Pro 2020 software (OriginLab).

#### D. Developing the machine learning kernel to analyze DCs migration

In this section, we explain the details of the data manipulation and the machine learning (ML) kernel (**Fig. 2**) to analyze DC migration. To construct the ML kernel, we prepared two separate datasets: one for training and the other for testing. The input data were manipulated by pre-processing, filtering, and polishing. After pre-processing, we obtained trajectories of 301 (imDCs) and 348 (mDCs) for training and 460 (imDCs) and 371 (mDCs) for testing. The results and figures in the manuscript were obtained from the testing dataset. The results from the training dataset are shown in **Figs. 2c**, **Extended data Fig. 2**, and **Extended data Fig. 3**.

Our ML kernel was designed to analyze complex and heterogeneous cellular motility. To analyze dynamic heterogeneity in a single DC migration, we segmented the trajectory data into 1 h long tracks and obtained a total of 8787 tracks from the testing dataset. For the training dataset, we prepared 5741 segmented tracks. From the segmented tracks, we extracted five features and applied two independent machine learning algorithms to determine distinct motility patterns that appeared during migration, regardless of cell maturation.

##### 1) Data pre-processing

**Data polishing.** We obtained the following raw trajectories of DC migration:348 (imDCs, training dataset), 383 (mDCs, training dataset), 449 (imDCs, testing dataset), and 350 (mDCs, testing dataset). Among them, a few raw trajectories were not tracked perfectly, and as a result, a few data points were absent in the trajectories. Therefore, we polished such trajectories. The number of imperfect trajectories (fraction) was identified as 16 (4.5%, imDC training dataset), 20 (5.22%, mDCs training dataset), 26 (5.79%, imDC testing dataset), and 20 (5.71%, mDCs testing dataset). The missing data points were attributed to tracking errors that mainly arise from cell-to-cell interactions during attachment between two distinct cells, and partially from the finite field-of-view, which results in the incomplete observation of cell movement near the edge of the observation window. Thus, we manually analyzed the trajectories using the following rule: First, we chose an arbitrary time window (≤10 min) and applied the spline interpolation provided in the Python library "scipy" package to the raw data[6]. If the untracked period was ≥10 min, we split the trajectory into two before and after attachment. Splitting an imperfectly tracked trajectory into two independent short trajectories does not seriously alter the statistics and the detection of migration patterns of single trajectories because this portion is approximately 4%–6% of the trajectory data. We also tested the interval of missing points, such as 20 min or 30 min time windows; however, the results shown in **Fig. 3** did not differ qualitatively. If we consider three dynamical modes, the statistical properties of the modes are robust against the interval of missing points owing to the relatively few imperfect trajectories obtained.

**Data filtering.** After data polishing, we removed inappropriate track samples in our ML analysis that fail to satisfy any of the following three criteria: Mean track speed ($\langle V_D \rangle_t \geq 1.5\ \mu\text{m/min}$), track duration ($T_{obs} \geq 60\ \text{min}$), and maximal displacement from the starting position ($\max|\vec{R}| \geq 20\ \mu\text{m}$)[3,7].

##### 2) Feature engineering

For the ML analysis, we calculated the five features shown below from the 1 h long segmented tracks. As explained in the main text, the five features are physical observables that enable quantification of the motility propensity and directionality of DC migration. The mathematical details of these features are as follows:

**(1) Radius of gyration.** To measure the second moments of positional fluctuations, we calculated the gyration tensor, $\mathbb{R}_{ij}$, which is given by

$$\mathbb{R}_{ij} = \frac{1}{N^2}\sum_{t_l=1}^{N}\sum_{t_m=1}^{N}(r_i(t_l) - \langle r_i \rangle)(r_j(t_m) - \langle r_j \rangle). \tag{1}$$

Here, the symbol $\langle O \rangle$ denotes time-averaging, and the subscripts $i, j$ represent the position coordinates $x, y$, respectively, and $l, m$ denote the time index runs from one to $N$ where $N$ is the trajectory length ($50 \leq N \leq 60$). The radius of gyration is obtained from the gyration tensor in Eq. (1) as follows:

$$R_g \equiv \sqrt{Tr\,\mathbb{R}}\,. \tag{2}$$

Because the $\langle r_i \rangle$ is the center of mass coordinate, $R_g$ represents the average spreading along the coordinates $x$ and $y$. Therefore, this quantity can be regarded as the spreading area per time unit.

**(2) Asphericity.** From Eq. (1), we can calculate the spreading of the migration trajectory along its principal axis. This shape-related property, called asphericity $A$, can be defined as

$$A \equiv \frac{(\lambda_1 - \lambda_2)^2}{(\lambda_1 + \lambda_2)^2} \tag{3}$$

where the $\lambda_{1,2}$ is the eigenvalue of the gyration tensor $\mathbb{R}_{ij}$. The asphericity is normalized in $[0,1]$ such that the two limiting values represent a perfect circular shape for zero and a straight-line shape for unity.

**(3) Energy consumption.** The degree of spreading in (2) can be related to the total kinetic energy spent over the entire migration event. To measure the relationship between the spreading and energy consumption, we evaluated the average kinetic energy as follows:

$$E \equiv \frac{1}{N-1} \sum_{i=0}^{N-1} V_D(t_n)^2. \tag{4}$$

Here, $V_D(t_i) = \frac{\sqrt{\left(r_x(t_{n+1}) - r_x(t_n)\right)^2 + \left(r_y(t_{n+1}) - r_y(t_n)\right)^2}}{(\mathrm{t}_{n+1} - t_n)}$ is the average migration speed at $t = t_n$ over 1 min.

**(4) End-to-end distance.** In addition to fast-spreading and high energy consumption, the directional propensity of migration can also affect the degree of overall spreading. We measured this effect in terms of the end-to-end distance.

$$R_{ete} = \sqrt{\left(r_x(t_N) - r_x(t_0)\right)^2 + \left(r_y(t_N) - r_y(t_0)\right)^2}. \tag{5}$$

Note that $R_{ete}$ is not the only time-averaged quantity among the features considered.

**(5) Variance of turning angles.** To measure the directional persistence, we have added a feature indicating the variance of turning angles. The $\theta$ is the angle between $\vec{D}(t_{n+1})$ and $\vec{D}(t_n)$ in the range $[-\pi, \pi)$, with a positive value in the counterclockwise direction where $\vec{D}(t_n; \Delta t) = \vec{r}(t_n + \Delta t) - \vec{r}(t_n)$. Using the segmented tracks, we calculated the variance of the turning angles as follows:

$$Var[\theta] \equiv \frac{1}{N-2} \sum_{i=1}^{N-2} (\theta_i - \langle \theta \rangle)^2 \tag{6}$$

##### 3) Training via cross-validation of two ML methods

With the five extracted features from the segmented trajectories, we constructed the input data of size 5741×5 for training the machine. First, we applied K-means unsupervised clustering to the input data to examine the number of distinct motility patterns of DCs[8,9]. Next, we used the results from K-means clustering as pseudo-labels and cross-validated these using the supervised learning algorithm XGBOOST[10]. The latter was trained with a subset of the input data (2000 trajectories) and we predicted the label for unknown trajectories in the segment pool with a 99.6% accuracy.

#### E. Identification of three distinct modes in DC migration

The trained XGBOOST algorithm was used to analyze unsegmented trajectories, and decipher the migration pattern in the 1 h interval time windows along the trajectory (**Fig. 2d**). We validated the ML method and its output by demonstrating that pattern classification is robust to variations in feature selection and their possible combinations.

##### 1) Number of distinct groups

We examined the average silhouette score to determine the number of distinct clusters in the migration patterns. The silhouette score has been widely used to infer the number of clusters in ML algorithms[11]. With the given cluster numbers as a hyperparameter, we measured the mean distance of a given data point within the cluster to which it belongs. For data point *i*, the mean distance within the cluster is estimated as $D_W(i) = \frac{1}{|C_I|}\sum_{j\in C_I} d(i,j)$, where $d(i,j)$ represents the Euclidian distance between the $i$ and $j$ data points in the cluster $C_I$, and $|C_I|$ is the size of the cluster. We estimated the dissimilarity of the data point $i$ by measuring the minimum distance between the data point and the other clusters, which is defined as $D_D(i) = \min_{\{C_J\}} \frac{1}{|C_J|}\sum_{j\in C_J} d(i,j)$, where $i \in C_I$, $j \in C_J$, and $\{C_J\}$ is the set of all neighboring clusters for $J \neq I$. The silhouette score for data point $i$ is defined as $s(i) = \frac{D_D(i)-D_W(i)}{\max[D_W(i),D_D(i)]}$. Thus, the average silhouette score lies between $[-1,1]$. The limit of 1 describes the situation in which the sample (i.e., cluster) is well separated from neighboring clusters.

In **Fig. 2c**, we show that the silhouette score reached the maximum when the number of distinct clusters was three if we used all five features introduced in our ML analysis. To test the robustness of the results, we performed ML analysis with various combinations of input features. For this test, we used a training dataset of 5741 segmented trajectories. In **Extended data Fig. 2**, we evaluated the silhouette scores for five distinct combinations of the input features. For all cases, we confirmed that the DC migration data most likely consisted of three dynamic modes.

##### 2) Statistical characteristics of the three distinct migration groups

We evaluated the statistical properties of the migration trajectories in each group with feature importance and the distribution of the five features (**Extended data Fig. 3**). **Extended data Fig. 3a** shows the feature importance provided by the XGBOOST algorithm. Asphericity, $A$, is the most important feature among the five when determining the group assignment. Here, we temporarily refer to the three groups as I, II, and III based on their population size.

The most dominant feature, $A$ plays a critical role in differentiating group I from groups II and III (**Extended data Fig. 3b**). The features, $R_g$, $R_{ete}$, and $E$, are effective in distinguishing Group II and Group III (**Extended data Fig. 3c, d, e**). These features quantify DC migration motility. In terms of $R_g$, $R_{ete}$, and $E$, groups I and II exhibited almost indistinguishable "slow" motility patterns, whereas group III with high motility was different from the other two groups. The variance of the turning angles $Var[\theta]$ had a minor contribution, and has a similar unimodal profile for all three groups (**Extended data Fig. 3f**). In **Table 1**, we have summarized a few important dynamic features of groups I, II, and III. Based on these properties, we assigned the dynamic modes of groups I, II, and III to slow-diffusive (SD), slow-persist (SP), and fast-persist (FP) phenotypes, respectively.

#### F. Analysis of dynamic properties for the three migration modes

We investigated the dynamic properties of the SD, SP, and FP modes based on commonly used statistical observables including mean-squared displacements (MSDs) and displacement probability density functions (PDFs). We also examined observables such as the turning angle heat map and density map of the phase space, $(V_n, \Delta\theta_n)$. Additionally, we investigated the zigzag-like patterns of DC migration in detail.

##### 1) Mean-squared displacements (MSDs)

From a single trajectory, we calculated the MSD as a function of the lag time $\Delta t$ using the following definition:

$$MSD = \frac{1}{T-\Delta t}\int_0^{T-\Delta t}[r(t+\Delta t)-r(t)]^2 dt \propto \Delta t^{\alpha} \ . \tag{7}$$

MSD often increases with $\Delta t$ in a power-law form. The power-law exponent $\alpha$ is called the anomalous exponent and provides dynamic information on the diffusion process: 1) $\alpha = 1$: where the diffusion is normal. 2) $0 < \alpha < 1$: diffusion is subdiffusive and consists of antipersistent random walks. 3) $1 < \alpha < 2$: Diffusion is super-diffusive and consists of persistent random walks. 4) $\alpha = 2$: The diffusion is ballistic such that the cell (or particle) moves at a constant velocity.

In terms of the anomalous exponent, DCs in the SD mode are sub-diffusive, and DCs in the SP and FP modes are super-diffusive. The plot indicates that cell migration dynamics change over time. The measured values of $\alpha$ in the short- and long-time regimes differed for all three dynamic modes. Compared with the averaged MSD, the individual MSD curves showed a broad scatter for the SD and SP modes (**Extended data Fig. 4**, gray lines in the upper plots), indicating the prevalence of substantial cell-to-cell variation in the SD and SP migrations. We quantified the (cell-to-cell) dynamic heterogeneity in terms of $P(\alpha)$ (**Extended data Fig. 4b**). The corresponding distribution for the SD mode showed the most spread from its peak value. Notably, the peak value of $P(\alpha)$ was approximately 0.25, whereas $\alpha$ for the averaged MSD was approximately 0.94. The result indicates that most of the individual SD-mode DCs moved more persistently than expected based on the averaged quantity. $P(\alpha)$ for the SP mode was also widely spread from its peak value. A few individual DCs exhibited subdiffusive motion, even though the averaged MSD indicates superdiffusive movement with $\alpha \approx 1.1 - 1.5$.

Compared to the SD and SP modes, FP-mode migration is relatively homogeneous. The $P(\alpha)$ was narrowly distributed around the peak value. A notable feature of the FP mode is that a few MSD curves show oscillatory behavior. We determined that such patterns can occur if the trajectory has a closed ellipse form in two-dimensional space. This occurs when a DC has positional fluctuations without meaningful movement comparable to the linear cell size. Although masked in the MSD plot, similar oscillating MSDs were found in the SD and SP modes.

##### 2) Displacement probability density functions (PDFs)

The displacement PDF is the Van Hove self-correlation function $P(x|\Delta t)$ where $x = x(t+\Delta t) - x(t)$ is the ($x$ -component) displacement over a given lag time $\Delta t$. As shown in **Extended data Fig. 5a,** we plotted the displacement PDFs at several lag times for the three modes. For all migration modes, the PDFs were neither Gaussian nor followed a power law. The profile of $P(x|\Delta t)$ provides additional information on migration dynamics. The non-Gaussianity indicates that the SD and SP modes cannot be governed by Gaussian-based anomalous diffusion models, such as fractional and scaled Brownian motion[12].

The exponential-like tail and cusp at the center in PDFs strongly suggests that DC migration is temporally heterogeneous and shows cell-to-cell heterogeneity. The fluctuation in the instantaneous diffusivity of DC movement and its wide distribution may result in the cusp and the exponential-like tail, as reported in studies using the fluctuating diffusivity model[13,14].

Interestingly, PDFs at different lag times can collapse onto a master curve if $x$ is re-scaled to $\frac{x}{\sqrt{<x^2>}}$ (**Extended data Fig. 5b**). We attempted to fit the master curves with nonlinear least-squares minimization ("lmfit" package for Python[15] via a stretched exponential function. See the caption for further information on the fitting).

##### 3) Turning angle heat map

The turning angle $\theta(\Delta t)$ refers to the angle between two consecutive displacement vectors. The displacement vector over a given lag time $\Delta t$ is defined as $\vec{D}(t_n;\Delta t) = \vec{r}(t_n + \Delta t) - \vec{r}(t_n)$ and $t_n = n\Delta t$. The turning angle was calculated as follows[18]:

$$\theta_n(\Delta t) = \cos^{-1}\left(\frac{\vec{D}(t_{n+1};\Delta t)\cdot\vec{D}(t_n;\Delta t)}{\left|\vec{D}(t_{n+1};\Delta t)\right|\left|\vec{D}(t_n;\Delta t)\right|}\right) \tag{8}$$

where the $\theta \in [-\pi, \pi]$, and the sign is positive (or negative) in the counterclockwise (or clockwise) direction, respectively. Next, we plotted the heat map of the normalized distribution for turning angles $P(\theta;\Delta t)$ as a function of the lag time. The corresponding heat maps for the three migration modes are shown in **Fig. 3d**. **Extended data Fig. 6** shows a plot of the heat maps obtained from a single trajectory. For each dynamic mode, we show ten randomly chosen heat maps to confirm whether the pattern in the averaged heat maps (**Fig. 3d**) is indeed observed at the single-trajectory level.

##### 4) Density map of phase space

To examine the radial and angular motions together, we constructed the phase-space $(\Delta\theta_n, V_n)$ where $\Delta\theta_n = \theta_{n+1} - \theta_n$, and $V_n = \frac{(|\vec{D}(t_{n+1};\Delta t)| - |\vec{D}(t_n;\Delta t)|)}{\Delta t}$. In **Extended data Fig. 7**, we show scatter plots of $(\Delta\theta_n, V_n)$ for several lag times in the three modes. For the radial contribution, the SD mode was symmetric, whereas the other two modes were asymmetric. For the latter modes, $V_n$ is thicker on the positive side or shifted to the right, which indicates that the cell moved away from the origin with time.

For the angular part, the SP mode showed a thick shoulder ($\pm\pi$) and small peaks at $\theta = \pm 2\pi$ for $\Delta t \geq 3$ min. When $\Delta\theta_n$ has peaks at $\pm 2\pi$ with a nonzero radial velocity, the corresponding trajectory shows two opposite turning events sequentially. The shoulder structure at $\pm\pi$ with a nonzero radial velocity may also indicate a zigzag-like motion, which will be described in the next section. The FP mode is characterized by a unimodal peak near zero, which shows a curved or straight motion by maintaining the turning angle for the overall relaxation time scale.

##### 5) Analysis of zigzag pattens in the DC migration data

In a recent study[16], a zig-zag Lévy walk was proposed as a phenomenological motility model for DC migration. The zigzag Lévy walk is characterized by repeating persistent runs, where each run is composed of zigzag-like (i.e., left-right or right-left) random walks. Here, we systematically analyzed the DC trajectories to examine whether zigzag migration patterns were prevalent during the DC migration dynamics. Our analysis showed that the SP mode is a plausible candidate for defining empirically observed zigzag migration.

For the analysis, we constructed the phase space of successive turning angles $(\theta_n, \theta_{n+1})$, where the subscript $n$ runs from the start-time segment to the last one. We show the density maps of $(\theta_n, \theta_{n+1})$ at several lag times for the three dynamic modes (**Extended data Fig. 8**). The schematic in **Extended data Fig. 8b** shows the trajectory motifs corresponding to the nine specific dense spots in the density maps. From the density maps (**Extended data Fig. 8c**), we calculated a zigzag fraction, which is the ratio of the total number of scatters between the first and third quadrants of the density map as a function of the lag time (**Extended data Fig. 9**). Note that our zigzag fraction is distinguished from the zig-zag preference factor introduced in reference,[16] in that our turning angles are defined from the displacement vectors of a given lag time, whereas in reference.[16] the turning events are determined using specific criteria, and the turning angles were obtained from these specific turning events.

The density maps in **Extended data Fig. 8c** show a few distinct patterns depending on the lag time and migration mode. At the shortest lag time ($\Delta t = 1\ min$), a common feature for all three modes is that the phase densities are concentrated around the origin. This suggests that the directional persistence of migration movement is conserved in this short-time regime (**Extended data Fig. 8a, b**). This tendency is consistent with the fact that directional persistence is lost at $\Delta t = 3$ min (**Fig. 3d**).

In the SD mode, as lag times increase, the dense concentration in the phase density at the origin becomes dispersed. At $\Delta t \geq 3$ min, the density map became dense at the four corners $(\pm\pi, \pm\pi)$, indicating that there exists an abundant pool of two motifs: $(\pi, \pi)$ and $(-\pi, -\pi)$, representing a circular motif, and $(\pi, -\pi)$ and $(-\pi, \pi)$, indicating the presence of zigzag-like motifs (**Extended data Fig. 8b**). However, the SD mode is not a zigzag migration in the long run because zigzag movements do not result in a persistent run. The zigzag fraction (the ratio of the total number of scatters between the first and third quadrants in the density map) in the SD mode migration was always less than unity (**Extended data Fig. 9**).

The SP mode shows nine dense spots for $\Delta t \geq 1$ min. In addition to the four corners $(\pm\pi, \pm\pi)$ and the origin, a dense area exists at $(\pm\pi, 0)$ and $(0, \pm\pi)$. This corresponds to the thick shoulder structure observed in **Extended data Fig. 7**, and the motifs exhibit a zig-zag motion (**Extended data Fig. 8b**). Because this phase space was obtained from two successive events, it does not directly represent zigzag migration. Nevertheless, the coexistence of a persistent motif, zigzag motif, and zigzag or zag motif suggests that the SP mode is a class of zigzag migrations. This view is also supported by the zigzag fraction pattern, which is larger than unity (**Extended data Fig. 9**).

For the FP mode, the phase density always includes a concentrated region at the origin, which can be expected due to its strong persistent movement over the entire observation period. The zigzag fraction increased to ~1.1, and subsequently decreased to saturation around unity (**Extended data Fig. 9**). It can be inferred that after a long relaxation time, the trajectory shape is curved due to strong directional persistence, indicating that the zigzag patterns are smeared (or show weak wiggling) owing to the large curves.

#### G. Molecular inhibition in the migration assay

##### 1) Inhibitor treatment

To study the role of myosin II and the Arp2/3 complex, the molecular inhibitors blebbistatin (20 μM, Sigma Aldrich) and CK666 (100 μM, Sigma Aldrich) were used during DC migration. Each inhibitor was mixed in an uncured gel (40 °C) before casting. After gel casting and curing, the gel confiner was incubated in complete medium with the same concentration of the inhibitor overnight. DMSO (0.1% (v/v); Sigma-Aldrich) was used as a negative control. Subsequently, the DCs located on the substrate were carefully covered with an inhibitor containing gel confiner. Cell motility assays were conducted after 1 h of incubation with covering by the gel confiner, similar to the approach described above.

**2) DC viability under inhibitor effect**

Ethidium homodimer (EthD-1; Thermo Fisher Scientific) was used to evaluate DC viability (**Extended data Fig. 10a**). After 24 h of incubation, DCs under gel confinement were stained with EthD-1 in complete medium (500 nM) in an incubator for 20 min. Fluorescent stained dead cells were counted manually using epifluorescence microscopy with a 20× Plan Apo lens (Nikon TiE). Cell survival rate was calculated as the number of stained cells per the number of whole cells in the field of view. More than 50 cells were analyzed for each experiment, and three independent experiments were performed.

#### References


1. Choi, Y., Kwon, J. E. & Cho, Y. K. Dendritic Cell Migration Is Tuned by Mechanical Stiffness of the Confining Space. *Cells* **10**, 3362 (2021).
2. Choi, Y., Sunkara, V., Lee, Y. & Cho, Y. K. Exhausted mature dendritic cells exhibit a slower and less persistent random motility but retain chemotaxis against CCL19. *Lab Chip* **22**, 377–386 (2022).
3. Vargas, P. *et al.* Innate control of actin nucleation determines two distinct migration behaviours in dendritic cells. *Nat Cell Biol* **18**, 43–53 (2016).
4. Normand, V., Lootens, D. L., Amici, E., Plucknett, K. P. & Aymard, P. New insight into agarose gel mechanical properties. *Biomacromolecules* **1**, 730–738 (2000).
5. Selinummi, J. *et al.* Bright Field Microscopy as an Alternative to Whole Cell Fluorescence in Automated Analysis of Macrophage Images. *PLOS ONE* **4**, e7497 (2009).
6. Virtanen, P. *et al.* SciPy 1.0: fundamental algorithms for scientific computing in Python. *Nat Methods* **17**, 261–272 (2020).
7. Stankevicins, L. *et al.* Deterministic actin waves as generators of cell polarization cues. *Proc Natl Acad Sci U A* **117**, 826–835 (2020).
8. MacQueen, J. Some methods for classification and analysis of multivariate observations. *Proc. Fifth Berkeley Symp. Math. Stat. Probab. Vol. 1 Stat.* **5.1**, 281–298 (1967).
9. Kanungo, T. *et al.* An efficient k-means clustering algorithm: analysis and implementation. *IEEE Trans. Pattern Anal. Mach. Intell.* **24**, 881–892 (2002).
10. Chen, T. & Guestrin, C. XGBoost. (2016).
11. Rousseeuw, P. J. Silhouettes - a Graphical Aid to the Interpretation and Validation of Cluster-Analysis. *J. Comput. Appl. Math.* **20**, 53–65 (1987).
12. Metzler, R., Jeon, J. H., Cherstvy, A. G. & Barkai, E. Anomalous diffusion models and their properties: non-stationarity, non-ergodicity, and ageing at the centenary of single particle tracking. *Phys Chem Chem Phys* **16**, 24128–24164 (2014).

13. Chubynsky, M. V. & Slater, G. W. 'Diffusing diffusivity': A model for anomalous and 'anomalous yet Brownian' diffusion. *Phys. Rev. Lett.* **113**, 098302 (2014).
14. Chechkin, A. V., Seno, F., Metzler, R. & Sokolov, I. M. Brownian yet Non-Gaussian Diffusion: From Superstatistics to Subordination of Diffusing Diffusivities. *Phys. Rev. X* **7**, 021002 (2017).
15. lmfit. https://lmfit.github.io/lmfit-py/intro.html - Google Search. https://www.google.com/search?q=Available+at%3A+https%3A%2F%2Flmfit.github.io%2Flmfit-py%2Fintro.html&oq=Available+at%3A+https%3A%2F%2Flmfit.github.io%2Flmfit-py%2Fintro.html&aqs=chrome..69i57.730j0j1&sourceid=chrome&ie=UTF-8.
16. Li, H. *et al.* Zigzag Generalized Levy Walk: the In Vivo Search Strategy of Immunocytes. *Theranostics* **5**, 1275–1290 (2015).

#### Supplementary Videos

**Supplementary Video 1**: Representative video of imDCs migrating under gel confinement. Lines represent the last 1 h trajectories and the color code indicates instantaneous speed. Time in hours:minutes. Background-subtraction was performed to increase the contrast of brightfield image.

**Supplementary Video 2**: Representative video of mDCs migrating under gel confinement. Lines represent the last 1 h trajectories and the color code indicates instantaneous speed. Time in hours:minutes. Background-subtraction was performed to increase the contrast of brightfield image.

**Supplementary Video 3**: Example of SD mode trajectory. A representative SD mode trajectory was collected in imDCs (represented by the blue line), and the line represents the last 1 h trajectory. Time in hours:minutes. Background-subtraction was performed to increase the contrast of brightfield image.

**Supplementary Video 4**: Example of SP mode trajectory. A representative SP mode trajectory was collected in imDCs (represented by the green line), and the line represents the last 1 h trajectory. Time in hours:minutes. Background-subtraction was performed to increase the contrast of brightfield image.

**Supplementary Video 5**: Example of FP mode trajectory. A representative FP mode trajectory was collected in mDCs (represented by the pink line), and the line represents the last 1 h trajectory. Time in hours:minutes. Background-subtraction was performed to increase the contrast of brightfield image.

**Supplementary Video 6**: imDC motility assigned to a machine-defined mode. Lines represent the last 1 h trajectories and the color code indicates the machine-defined mode (SD: blue, SP: green, FP: pink). Time in hours:minutes. Background-subtraction was performed to increase the contrast of brightfield image.

**Supplementary Video 7**: mDC motility assigned to the machine-defined mode. Lines represent the last 1 h trajectories and the color code indicates the machine-defined mode (SD: blue, SP: green, FP: pink). Time in hours:minutes. Background-subtraction was performed to increase the contrast of brightfield image.

**Supplementary Video 8**: imDC motility after 20 μM blebbistatin treatment. Lines represent the last 1 h trajectories and the color code indicates the machine-defined mode (SD: blue, SP: green, FP: pink). Time in hours:minutes. Background-subtraction was performed to increase the contrast of brightfield image.

**Supplementary Video 9**: imDC motility after 100 μM CK666 treatment. Lines represent the last 1 h trajectories and the color code indicates the machine-defined mode (SD: blue, SP: green, FP: pink). Time in hours:minutes. Background-subtraction was performed to increase the contrast of brightfield image.

**Supplementary Video 10**: mDC motility after 20 μM blebbistatin treatment. Lines represent the last 1 h trajectories and the color code indicates the machine-defined mode (SD: blue, SP: green, FP: pink). Time in hours:minutes. Background-subtraction was performed to increase the contrast of brightfield image.

**Supplementary Video 11**: mDC motility after 100 μM CK666 treatment. Lines represent the last 1 h trajectories and the color code indicates the machine-defined mode (SD: blue, SP: green, FP: pink). Time in hours:minutes. Background-subtraction was performed to increase the contrast of brightfield image.

**Supplementary Video 12**: Representative cyclic mode transition in imDCs. Lines represent the last 1 h trajectories and the color code indicates the machine-defined mode (SD: blue, SP: green, FP: pink). Time in hours:minutes. Background-subtraction was performed to increase the contrast of brightfield image.
